# Length Biases in Single-Cell RNA Sequencing of pre-mRNA

**DOI:** 10.1101/2021.07.30.454514

**Authors:** Gennady Gorin, Lior Pachter

## Abstract

Single-molecule pre-mRNA and mRNA sequencing data can be modeled and analyzed using the Markov chain formalism to yield genome-wide insights into transcription. However, quantitative inference with such data requires careful assessment and understanding of noise sources. We find that long pre-mRNA transcripts are over-represented in sequencing data, and explore the mechanistic implications. A biological explanation for this phenomenon within our modeling framework requires unrealistic transcriptional parameters, leading us to posit a length-based model of capture bias. We provide solutions for this model, and use them to find concordant and mechanistically plausible parameter trends across data from multiple single-cell RNA-seq experiments in several species.

## 1 Background

The development of increasingly accurate and quantitative single-cell RNA sequencing (scRNA-seq) makes it increasingly tractable to fit single-molecule data to models of the RNA life cycle, thus facilitating a *mechanistic* view of genome-wide transcriptional regulation. Specifically, protocols with cell barcodes and unique molecular identifiers (UMIs) allow for parameterization of discrete probabilistic models, whereby it is natural to interpret the contents of cells as draws from distributions over the non-negative integers.

A standard framework for such modeling is the Chemical Master Equation (CME), which models the molecular contents of cells via Markov chains that traverse a discrete state space of RNA counts [1–3]. To fit biophysical parameters (the “inverse” problem of inference), one must solve the CME (the “forward” problem of prediction). This workflow requires computationally facile solutions that can be applied to thousands of genes. In mammalian and bacterial systems, the specific form of the CME is based on a *random telegraph* model of gene regulation, which describes a single gene locus that randomly switches between active and inactive states [1]. A common simplification, supported by genome-wide fluorescence studies [4], treats the active state’s duration as vanishingly small: mRNA is produced in geometrically-distributed bursts that arrive according to a Poisson process. This model can be extended to describe rather general *downstream* processes of splicing, degradation, and translation. We focus on newly available data with spliced and unspliced mRNA, which can be fit to an analytically tractable bursting model [5,6].

Within this classical framework, there are two substantial challenges that must be overcome in order to perform inference with scRNA-seq data: first, sequencing is probabilistic, with some molecules inevitably unobserved in the process. Second, conventional “top-down” methods for scRNA-seq data processing used to correct for sampling artifacts, such as hyperparametrized imputation, normalization, and smoothing [7], are incompatible with a discrete “bottom-up” picture of the underlying biophysics. A variety of approaches have been adopted to address these issues. Certain methods abandon the CME altogether, instead applying moment-based models to smoothed and normalized data [8,9]; this approach may introduce distortions of an unknown magnitude, and can result in loss of statistical power. Other approaches do not correct for sampling [10, 11], with resultant batch effects of an unknown magnitude persisting in the data. Thus, treating both biological and technical stochasticity remains a significant lacuna in single-cell transcriptional models, with no satisfactory and rigorous solutions.

## 2 Length bias in measured pre-mRNA expression

We begin by exploring the biophysical interpretability of scRNA-seq data in light of the length bias seen in pre-mRNA expression. Such analysis is crucial for understanding approaches such as RNA velocity [8,12]. In some datasets, average spliced mRNA counts do not seem to show a length dependence (Fig. 1a), consistently with previous studies of UMI-based protocols [13]. On the other hand, unspliced mRNA counts strongly correlate with gene length (Fig. 1b). This prompts us to investigate whether this discrepancy has biological origins, and raises questions about the consequences of ignoring this bias.

**Figure 1:**
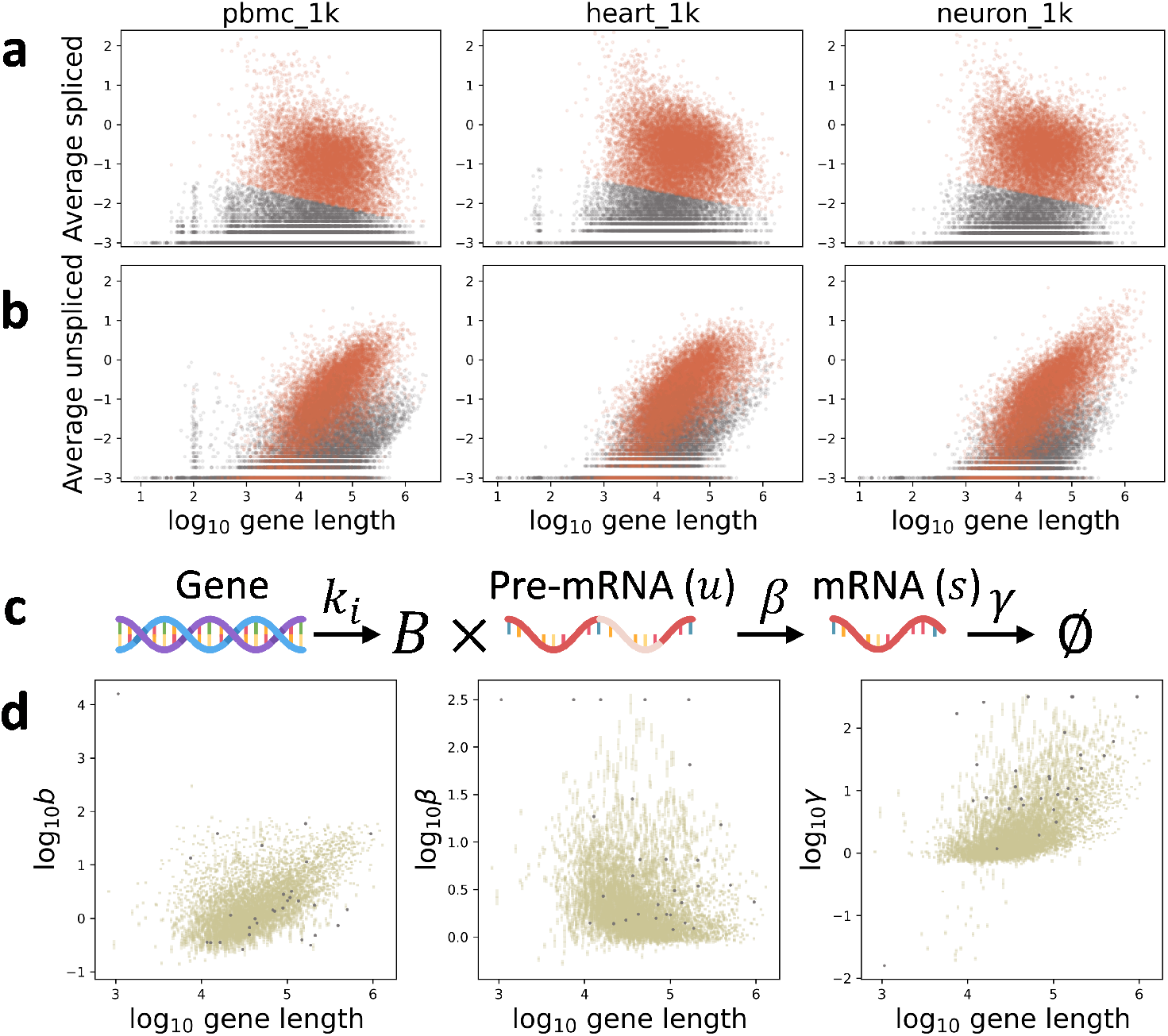
**a.** Length dependence of average spliced mRNA observations in three datasets (orange: high-expression gene cluster; gray: discarded low-expression cluster). **b.** Length dependence of average unspliced mRNA observations in same datasets (colors as in **a**). **c.** Model of transcriptional physiology. **d.** Transcriptional parameter estimates without a stochastic model of sequencing demonstrate pervasive length-dependent trends (gold: lower bounds on 99% confidence intervals; gray: fits rejected by statistical testing).

### 2.1 Stochastic analysis produces implausible parameter estimates

At first glance, a naïve analysis using a conventional [4] stochastic transcriptional model (Fig. 1c as described in Section 3.2) yields reasonable results (Fig. 1d). However, comparison with previous transcriptome-wide analyses prompts questions regarding the plausibility of these findings and the model’s ability to represent scRNA-seq data.

We find that the inferred burst size increases with transcript length, in stark contrast with the previously observed modest inverse relationship [14]. The degradation rate, normalized to burst frequency, displays a similar positive trend. Previous studies found little to no gene length effect on burst frequency [14] and no effect on the rate of mRNA degradation [15]. The latter is primarily controlled by open reading frame features rather than the length of the source gene. The decreasing rates of splicing are more challenging to analyze: the splicing timescales given in the literature vary over several orders of magnitude depending on system and technology [16]. However, several factors suggest that length-based effects should be minimal: co-transcriptional splicing is expected to be ubiquitous in mammalian cells [17,18], and widely varying intron sizes have little impact on splicing time in direct comparison using a single protocol [19]. Therefore, the parameter estimates are discordant with previously reported results.

Aside from empirical data, there are theoretical reasons to question these results: there is little reason any of these parameters should “know” about the gene length. Although transcription of longer genes should ostensibly take longer, it seems reasonable to posit that the burst frequency and size should be governed by the *promoter* rather than any downstream elements. Similarly, the splicing rate is likely governed by spliceosome kinetics at individual introns, which is a local, rather than gene-wide, effect. Finally, the transcript that is degraded is the terminal isoform, whose length is only weakly related to the length of the original parent transcript.

In summary, the observed UMI counts of spliced transcripts cannot be plausibly treated the same way as those of unspliced transcripts. Such a simplification is incompatible with empirical evidence and currently accepted models.

## 3 A technical noise model

In the current section, we motivate, solve, and apply a *stochastic* model of sequencing that addresses technical artifact to scRNA-seq data. First, we describe the primary probability objects in Section 3.1. We use the CME framework to derive the model from a microscopic Markov description of transcription in Section 3.2. Finally, we report the model solution in Section 3.3, and fully describe the derivation in Section S1.1.

In brief, we build a model that explicitly incorporates the stochastic sequencing steps taking place in fixed media (Fig. 2a). Consistently with previous work on modeling pre-mRNA [8], we assume that the library construction step in the 10X sequencing workflow [20] includes molecules that have been captured at off-target binding sites. We posit that unspliced mRNA are primarily captured at internal poly(A) stretches, whereas spliced mRNA are captured at the poly(A) tail. To quantitatively model this effect, we introduce the concept of UMI “false positives”: if a molecule has sufficiently many poly(A) sites, it is likely to be captured and reverse-transcribed multiple times. As a first-order approximation, we model this bias as a length-dependent capture rate. Thus, each molecule in a cell gives rise to a Poisson distribution of cDNA. The downstream sequencing and alignment steps are treated as binomial sampling from the cDNA distribution.

**Figure 2:**
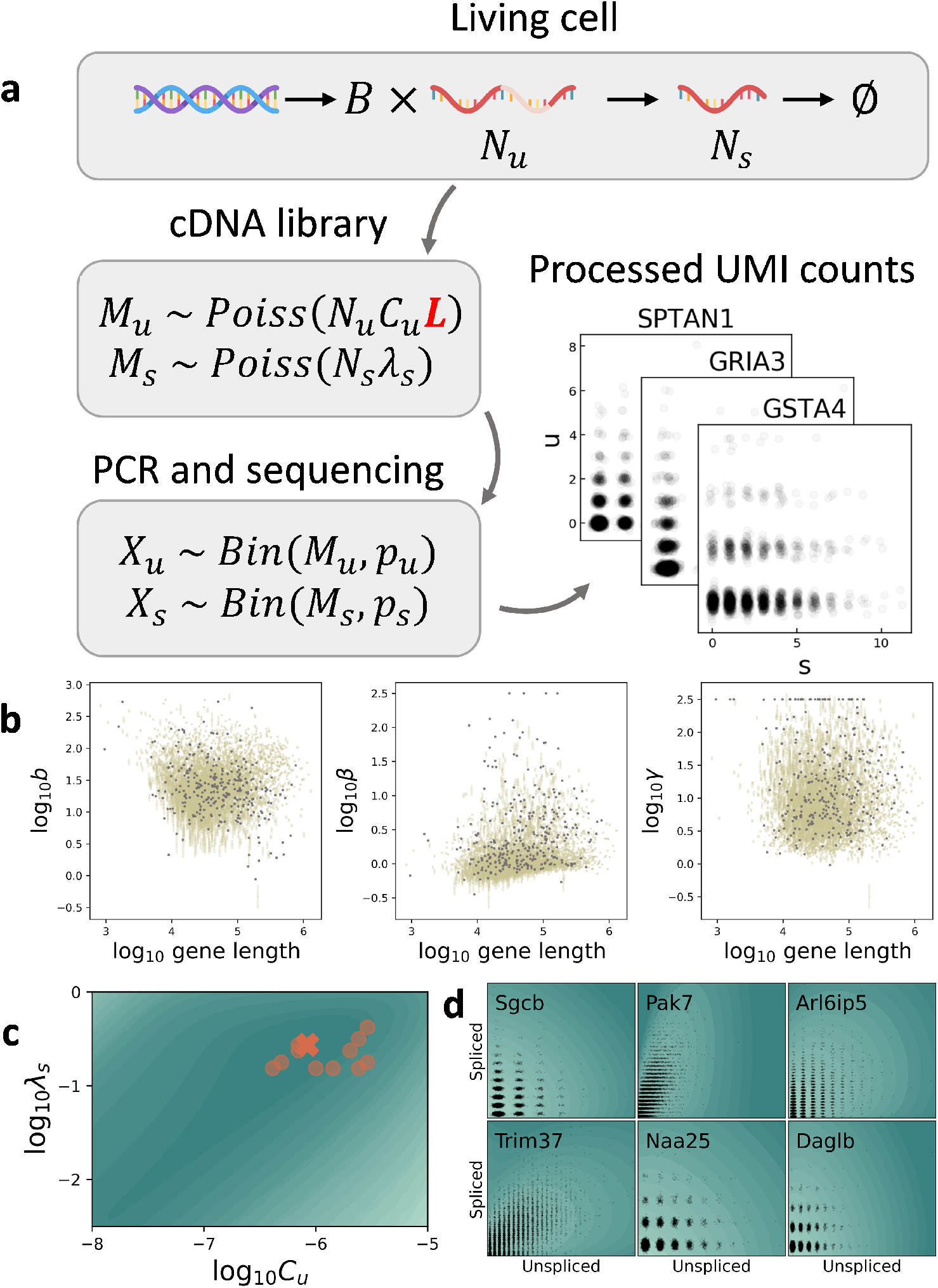
**a.** The integrated stochastic model of transcription and sequencing, with length dependence of the library construction step indicated in red. **b.** Inferred transcriptional parameters do not appear to have strong length dependence (gold: lower bounds on 99% confidence intervals; gray: fits rejected by statistical testing). **c.** The sampling parameter likelihood landscape shows a single optimum (dark: lower, light: higher total Kullback-Leibler divergence between fit and data; orange cross: optimal sampling parameter fit for the displayed landscape; orange points: optimal sampling parameter fits for other analyzed datasets). **d.** The parameter fitting procedure successfully recapitulates empirical observations (dark: lower, light: higher log probability mass; black points: raw data UMI counts).

### 3.1 Probability distributions

A probability density function (PDF) associated with a continuous-valued random variable is denoted by *f* (·;·). A probability mass function (PMF) associated with a discrete-valued random variable *X* is denoted by *P*(*X* = *x*;·) or *P*(*x*;·). Analogously, the joint multivariate PMF values are given by *P* (*X*_1_ = *x*_1_, *X*_2_ = *x*_2_,…;·) or *P* (*x*_1_, *x*_2_,…;·). The parameter arguments are elided throughout the rest of the paper for simplicity of notation. The PMF of a nonnegative-valued discrete random variable can be equivalently expressed as the continuous probability generating function (PGF), 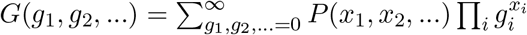. The multivariate PGF is given by 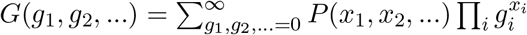.

The following distributions arise in the problem:

- Exponential: if *X* ~ *Exp*(λ), *f*(*x*; λ) = λ*e*^−λ*x*^.
- Geometric: if *X* ~ *Geom*(*p*), *P*(*X* = *k*;*p*) = (1 – *p*)^*k*^*p*, where *p* ∈ (0,1] and 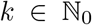. The geometric distribution is well-known to arise in the short-burst limit of the two-state transcription model [21]. The corresponding PGF is 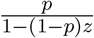.
- Negative binomial (NB): if *X* ~ *NegBin*(*r,p*), 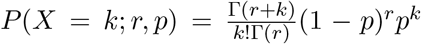, where *p* ∈ [0,1] and *r* > 0. We note that MATLAB and the NumPy library take the opposite convention, with a 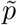 parameter defined as 1 – *p*. The corresponding PGF is 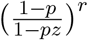.
- Poisson: if *X* ~ *Poisson*(λ), 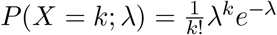. The corresponding PGF is *e*^λ(*z*−1)^.
- Binomial: if *X* ~ *Bin*(*n,p*), 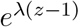. The corresponding PGF is [(1 – *p*) + *pz*]^*n*^.
- Bernoulli: if *X* ~ *Bernoulli*(*p*), *P*(*X* = *k*;*p*) = *p*^*k*^(1 – *p*)^1–*k*^. The corresponding PGF is (1 – *p*) + *pz*. The Bernoulli distribution is a degenerate case of the binomial distribution, with *n* = 1.

### 3.2 Model definition

We summarize the model-specific quantities, distributions, and random variables in Table 1, and define the relevant parameters in Table 2. In the living cell, we model a two-stage birth-death process coupled to a bursting promoter:

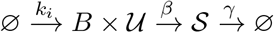

**Table 1:** Theoretical statistical variables

| Variable | Definition |
| --- | --- |
| $z$ | Molecular species, either unspliced $u$ or spliced $s$ |
| $n_z$ | Number of *in vivo* mRNA (species $z$) |
| $N_z$ | Random variable describing number of *in vivo* mRNA (species $z$) |
| $m_z$ | Number of *in vitro* UMIs for cDNA (species $z$) |
| $M_z$ | Random variable describing number of *in vitro* cDNA UMIs (species $z$) |
| $x_z$ | Number of *in silico* sequenced and identified UMIs (species $z$) |
| $X_z$ | Random variable describing number of *in silico* UMIs (species $z$) |
| $P(n_u, n_s)$ | Joint PMF of unspliced and spliced mRNA |
| $G(g_u, g_s)$ | PGF of $P(n_u, n_s)$ |
| $P(m_u, m_s)$ | Joint PMF of unspliced and spliced cDNA UMI counts |
| $G_{1,z}(g_z)$ | Per-molecule PGF of cDNA capture process |
| $G_{M_u, M_s}(g_u, g_s)$ | PGF of $P(m_u, m_s)$ |
| $P(x_u, x_s)$ | Joint PMF of unspliced and spliced identified UMI counts |
| $G_{2,z}(g_z)$ | Per-molecule PGF of sequencing and identification process |
| $G_{X_u, X_s}(g_u, g_s) = H$ | PGF of $P(x_u, x_s)$ |

**Table 2:** System parameters

| Parameter | Definition |
| --- | --- |
| $z$ | Molecular species, either unspliced $u$ or spliced $s$ |
| $k_i$ | Burst frequency |
| $B$ | Stochastic burst size |
| $b$ | Expectation of $B$ |
| $\beta$ | Splicing rate |
| $\gamma$ | Degradation rate |
| $D_z$ | Poisson rate of per-molecule cDNA generation |
| $p_z$ | Bernoulli probability of per-molecule sequencing and identification |
| $\lambda_z$ | $D_z p_z$, aggregated Poisson rate of per-molecule *in silico* UMI generation |
| $L$ | Gene length |
| $C_u$ | Length dependence of $\lambda_u$, such that $\lambda_u = C_u L$ |

The bursts arrive with frequency *k_i_*, and generate *u*nspliced mRNA. The bursts have magnitude 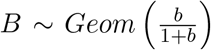, with expectation *b*. After a waiting time ~ *Exp*(*β*), the unspliced mRNA is converted to spliced mRNA. After another waiting time ~ *Exp*(*γ*), the spliced mRNA is degraded. We assume the system reaches its unique steady state; the existence of such a state is guaranteed by ergodicity. This in *vivo* stage produces an unobservable stationary distribution *P*(*N_u_* = *n_u_*, *N_s_* = *n_s_*), where *N_z_* is the random variable describing true physiological counts of molecule *z* and *n_z_* is the molecule count. The PGF of this distribution is defined as *G*(*g_u_*, *g_s_*).

After equilibration, cDNA library construction begins and all physiological processes halt due to cell fixation[20]. Due to the possibility of multiple priming, each molecule of mRNA produces *Poisson*(*D_z_*) molecules of cDNA. *D_u_* is presumed length-dependent and governed by internal priming, whereas *D_s_* is presumed length-independent and governed by poly(A) tail priming. This stage produces a distribution *P*(*M_u_* = *m_u_*, *M_s_* = *m_s_*), where *M_z_* is the random variable describing UMI counts of a gene product and *m_z_* is the cDNA number. The definition of the sampling procedure implies *M_z_*|*n_z_* ~ *Poisson*(*D_z_n_z_*). As we assume the species are sampled independently, we can represent the per-molecule PGF as *G*_1,*z*_(*g_z_*) = *e*^*D_z_*(*g_z_* − 1)^. The corresponding overall joint PGF is *G*_*M_u_*,*M_S_*_(*g_u_*, *g_s_*) = *G*(*G*_1,*u*_(*g_u_*),*G*_1,*s*_(*g*_s_)).

Finally, amplification and sequencing take place. Significantly, unlike the library construction, these are strictly *depleting* processes: we suppose they cannot generate new UMIs, but they *can* lead to loss of UMIs. We assume the PCR amplification and product fragmentation are not substantially biased from gene to gene; further, the downstream fragments do not retain length information. Nevertheless, the overall identifiability of unspliced and mature mRNA may be different. Therefore, we suppose that each *in vitro* cDNA UMI gives rise to *Bernoulli*(*p_z_*) ∈ {0,1} amplified, sequenced, and corrected *in silico* UMIs. The definition of the sampling procedure implies *X_z_*|*m_z_* ~ *Bin*(*m_z_*, *p_z_*). We can represent the per-molecule PGF as *G*_2,*z*_(*g_z_*) = (1 – *p_z_*) + *p_z_g_z_*. The corresponding overall joint PGF is *G*_*X_u_*, *X_s_*_(*g_u_*,*g_s_*) = *G*(*G*_1,*u*_(*G*_2,*u*_(*g_u_*)),*G*_1,*s*_(*G*_2,*s*_(*g_s_*))) ≔ *H*(*g_u_*, *g_s_*). The parameters *D_z_* and *p_z_* are not independently identifiable, leading us to define net sampling rates *λ_z_* ≔ *D_z_p_z_*.

We use a first-order model of length dependence λ_*u*_ = *C_u_L*, which implies that the rate of capture of any particular molecule scales directly with its length, acting as a proxy for the number of poly(A) stretches in the molecule. It is well-known that even short poly(A) sequences can be captured by the oligo(dT) primers used in sequencing [22], and the number of poly(A) sequences in a given gene is strongly correlated with length (Fig. S2). We do not directly consider the number of stretches, as the determination of appropriate length thresholds or weights is a distinct thermodynamics challenge. The spliced mRNA parameter λ_*s*_ is kept constant, modeling capture at the poly(A) tail.

### 3.3 Model solution

Per previous reports [5,6], the steady-state PGF for the joint distribution of unspliced and spliced mRNA is *G*(*g_u_*, *g_s_*) = *e*^*ϕ*(*v_u_*, *v_s_*)^, where *v_u_* = *g_u_* − 1, *v_s_* = *g_s_* − 1, 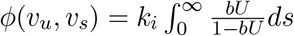, 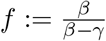, and *U* = *v_s_fe*^−γ*s*^ + [*v_u_* − *v_s_f*]*e*^−*βs*^. As derived above, the PGF of a distribution under two steps of independent sampling is *H*(*g_u_*, *g_s_*) = *G*(*G*_1,*u*_(*G*_2,*u*_(*g_u_*)), *G*_1,*s*_(*G*_2,*s*_(*g_s_*))), where *G_i,z_* is the PGF for sampling step *i* and species *z*. Using the model assumptions outlined above, the overall PGF *H*(*g_u_*, *g_s_*) = *G*(*e*^λ_*u*_(*g_u_* − 1)^, *e*^λ_*s*_(*g*_*s*_ − 1)^). The corresponding joint probability distribution *P*(*x_u_*, *x_s_*) is easily computed by evaluating *g_u_* and *g_s_* around the complex unit circle and performing an inverse Fourier transform.

The moments of the model can be calculated using the definition of the PGF. We define the relevant theoretical and empirical summary statistics in Table 3, and report the lower moments of the noise-free model and the full model in Table 4.

**Table 3:**
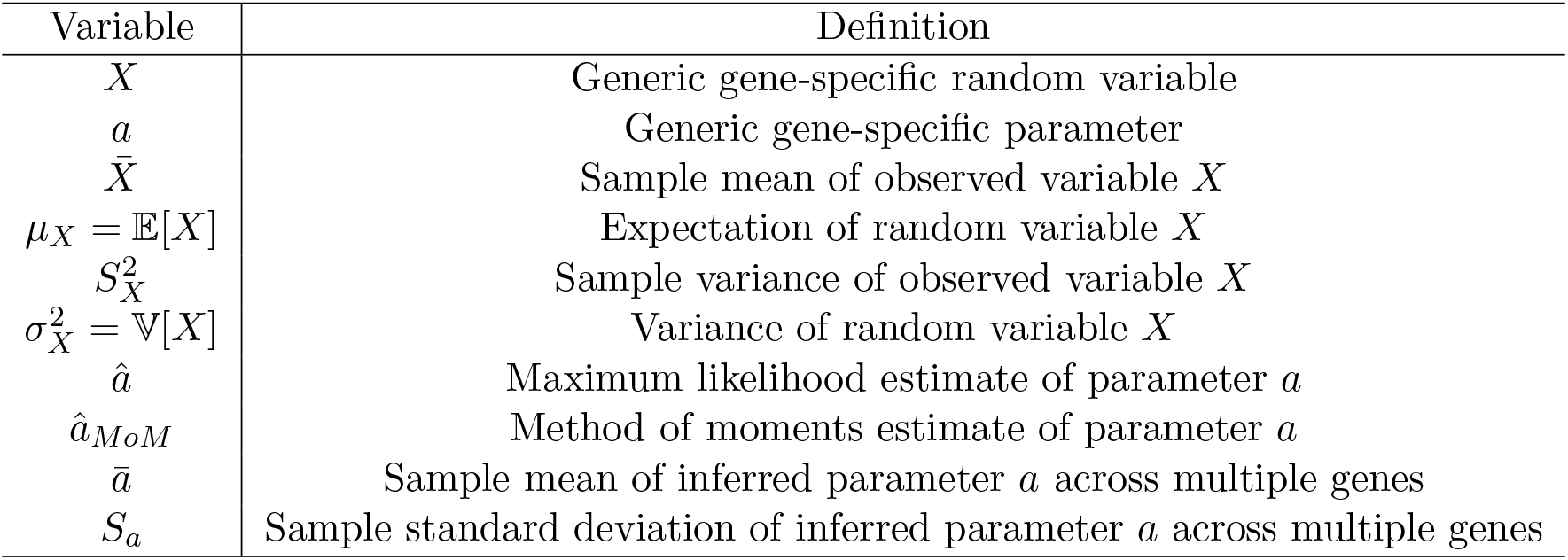
Summary statistics

| Variable | Definition |
| --- | --- |
| $X$ | Generic gene-specific random variable |
| $a$ | Generic gene-specific parameter |
| $\bar{X}$ | Sample mean of observed variable $X$ |
| $\mu_X = \mathbb{E}[X]$ | Expectation of random variable $X$ |
| $S_X^2$ | Sample variance of observed variable $X$ |
| $\sigma_X^2 = \mathbb{V}[X]$ | Variance of random variable $X$ |
| $\hat{a}$ | Maximum likelihood estimate of parameter $a$ |
| $\hat{a}_{MoM}$ | Method of moments estimate of parameter $a$ |
| $\bar{a}$ | Sample mean of inferred parameter $a$ across multiple genes |
| $S_a$ | Sample standard deviation of inferred parameter $a$ across multiple genes |

**Table 4:** Summary statistics

| Moment | Noise-free model | Technical noise model |
| --- | --- | --- |
| $\mu_u$ | $\frac{k_i b}{\beta}$ | $\frac{\lambda_u k_i b}{\beta}$ |
| $\mu_s$ | $\frac{k_i b}{\gamma}$ | $\frac{\lambda_s k_i b}{\gamma}$ |
| $\sigma_u^2 - \mu_u$ | $\mu_u b$ | $\mu_u \lambda_u (1 + b)$ |
| $\sigma_s^2 - \mu_s$ | $\mu_s \frac{b\beta}{\beta+\gamma}$ | $\mu_s \lambda_s \left(1 + \frac{b\beta}{\beta+\gamma}\right)$ |
| $\text{Cov}(u, s)$ | $\frac{k_i b^2}{\beta+\gamma}$ | $\frac{\lambda_u \lambda_s k_i b^2}{\beta+\gamma}$ |

## 4 Experimental data yield consistent parameter fits

### 4.1 Data processing and inference

The processing pipeline is summarized in Fig. S1. Briefly, we initialize a grid of sampling parameters, compute the conditional maximum likelihood estimates of the transcriptional parameters for each grid point, identify the global optimum, then analyze the parameter trends.

We collected *Ensembl* FASTQ files [23] corresponding to the full human and mouse genomes, computed gene lengths, and partitioned each gene’s sequence into a set of contiguous poly(A) sequences. These sequences were used to compute cumulative histograms of the number of poly(A) stretches.

We collected FASTQ produced by 10X scRNA-seq and processed them with the `kallisto|bustools` pipeline [12]. The twelve analyzed datasets are summarized in Table S1; eight were generated by 10X Genomics and four were generated for a mouse brain cell atlas study [24, 25]. Pseudoalignment and quantification produced a set of *loom* files. The numbers of cells and genes retained after filtering are reported in the “Cells” and “Genes Detected” columns. Genes without length annotations were discarded. As shown in Fig. S4, all datasets demonstrated the previously encountered (Fig. 1) expression bias. Genes appeared to fall into two categories, with two distinct linear trends: high-expression genes (orange) and low-expression genes (gray). We clustered [26] based on the spliced expression/length plot and discarded the low-expression genes, with effective resolution of the unspliced expression trends. As quantified in Table S2, the low-expression cluster was dominated by pseudogenes and non-coding genes with poor annotations (over 50% of the cluster in each dataset). We likewise removed genes considered too sparse (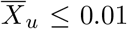, 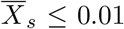, max *X_u_* ≤ 3, max *X_s_* ≤ 3) or too computationally intensive (max *X_u_* ≥ 350, max*X_s_* ≥ 350) to fit. The numbers of genes retained after filtering are reported in the “Genes Kept” column of Table S1.

The datasets lend themselves to interpretation as technical and biological replicates: for example, “10X 1k PBMC” and “10X 10k PBMC” correspond to two libraries produced from a single human blood cell sample, whereas “Allen B01” and “Allen A08” correspond to libraries produced from the brains of two distinct mice. Therefore, we split the twelve datasets into the four subsets reflected in Table S1 and identified the genes retained after filtering in all members of the subset; the size of the set is reported in the “Genes Overlap” column. Finally, we selected a random subset of these genes to fit; the number of genes is reported in the “Genes Selected” column. The names of genes rejected by clustering, as well as those analyzed using the workflow, are provided in Supplementary File 1.

Parameter estimation was performed by minimizing the Kullback-Leibler (KL) divergence between the data and the proposed PMFs. We computed each PMF using the solution reported in Section 3.3, evaluated over the data domain. The integral was approximated computed using order 60 Gaussian quadrature on *t* ∈ [0,10(*β*^−1^ + *γ*^−1^)], implemented using `scipy.integrate.fixed_quad`. The alternative adaptive procedure `scipy.integrate.quad_vec` is available, but requires an order of magnitude more time for evaluation. The incurred error was small enough to justify the approximation (no more than 10^−4^ total variation distance).

Fits to the noise-free model were performed by gradient descent initialized at the method of moments parameter estimates (Section S1.2.3), computed from the moments reported in Table 4. Identifying parameters of the technical noise model was somewhat more challenging: given *N* genes, the model has 3*N local* (gene-specific) and 2 *global* (genome-wide) parameters. To fit all parameters, we implemented a variant of the coordinate descent algorithm: we scanned over a grid of *C_u_* and λ_s_ and computed *conditional* maximum likelihood estimates *b*, *β*, *γ*|*C_u_*, λ_*s*_. For each tuple of (*C_u_*, λ_*s*_), the optimization routine was initialized at the method of moments estimates (Section S1.2.3). The global maximum likelihood estimate was computed by identifying the sampling parameter set with lowest total divergence. The scan and search domains are reported in Table S3; a 40 × 41 grid was used. The coordinate descent approach also permitted direct examination of the likelihood landscapes, and allowed us to confirm the existence of a unique sampling parameter optimum. Finally, we applied the chi-squared test with the Bonferroni correction to each optimal PMF, and discarded all genes with parameters within 0.01 of the search domain bounds to identify potential model misspecification problems.

To compute confidence intervals, we used local Gaussian approximation to the maximum likelihood estimate, justified by the relatively high number of samples (cells). For the noise-free model, we computed the Fisher information matrix (Hessian of the KL divergence) *I* at the maximum likelihood estimate 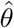, inverted it, and found lower bounds on the parameter standard deviations as the diagonal entries 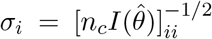, where *n_c_* denotes the number of cells. Given these standard deviations *σ_i_*, we used the z-score to estimate 99% confidence intervals as 2.576*σ_i_*. For the technical noise model, we determined the Fisher information matrix using an identical computation, holding the sampling parameters fixed. This procedure effectively yields a *conditional* posterior distribution that is necessarily an underestimate, because uncertainty in {*C_u_*,λ_*s*_} is unaccounted for, but consistent with the interpretation of *σ_i_* as a theoretical lower bound.

### 4.2 Results

Fitting transcriptional parameters using the noise-free model produced trends consistent with the fits described above (Fig. S5), and inconsistent with orthogonal empirical results (Section 2.1). Two other noise models without a sequencing length bias produced qualitatively identical results (Section S2). Fitting the Poisson technical noise model yielded transcriptional parameters (Fig. 2b, Fig. S6) with very weak length dependence. Therefore, we suggest that this integrated description of transcription and sequencing provides a more realistic and physically interpretable picture than available by considering the two sources of stochasticity separately.

All optima discovered by the coordinate scan procedure lie within the square log_10_*C_u_* = −6.0 ± 0.4 and log_10_λ_*s*_ = −0.6 ± 0.2. The KL divergence landscapes suggest that the datasets have unique optima and the model is appropriate (Fig. 2c, Fig. S7). Furthermore, empirical joint mRNA count histograms were consistent with the fits (Fig. 2d).

The inferred parameter distributions were consistently well-described by a log normal-inverse Gaussian law (Fig. 3a, Fig. S8), although the mechanistic import of this finding is unclear. We performed a set of technical replicates, fitting distinct libraries generated from the same organism, and biological replicates, fitting libraries from multiple organisms. The results (Fig. 3b-c, Fig. S9, S10) are consistent, with higher correlations among the technical replicates. We report the statistical summaries of maximum likelihood parameter fits, as well as the fractions of rejected genes, in Table S4.

**Figure 3:**
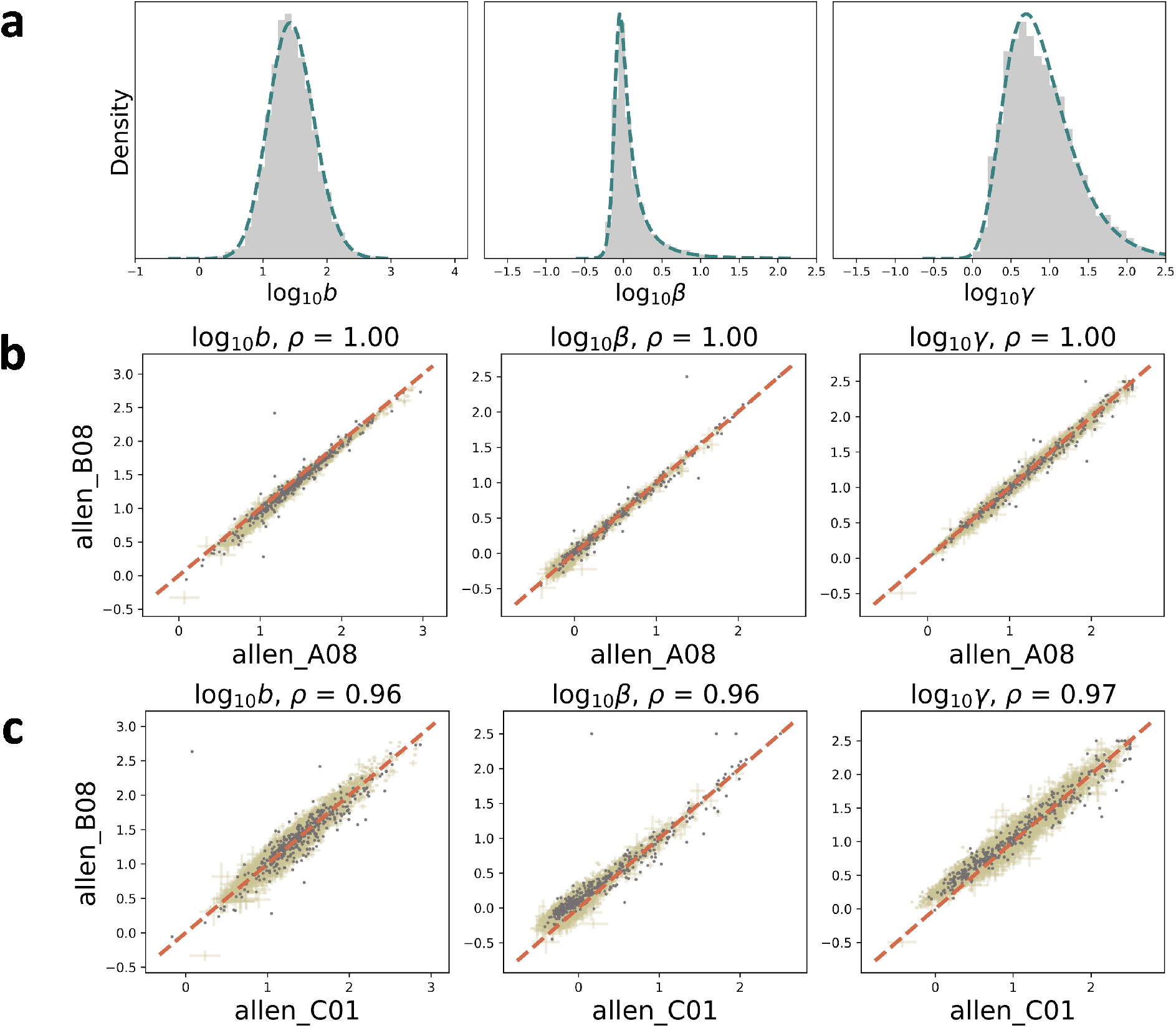
**a.** Inferred transcriptional parameter distributions (`allen_B08`; gray: histogram of 4696 genes retained after statistical testing; teal line: best fit to normal-inverse Gaussian distribution). **b.** Technical replicates show largely concordant inferred parameter values (orange dashed line: identity; gold: lower bounds on 99% confidence intervals; gray: fits rejected by statistical testing).**c.** Biological replicate parameter estimates are likewise concordant (colors as in **b**).

Using the same procedure, we compared whole-cell and nucleus-only libraries generated from mouse neurons. The nucleus-only fits had significantly worse performance (50.5% rejected by chi-squared, vs. 9.1%), and demonstrated a dramatic offset with respect to the whole-cell fits, particularly for the degradation parameter (Fig. S11). Comparison of successful fits to all datasets (Fig. S12) appears to confirm this effect. Average parameter values vary over roughly an order of magnitude; however, the nuclear RNA dataset has a rather higher location parameter.

Comparing parameter fits for a subset of genes observed in heart and neuron datasets, we observed much weaker correlations (Fig. S13). This effect is concordant with expectations, as different tissues have different gene expression patterns. In the case of splicing and degradation rate, we observed pervasive offsets (on the order of 10^0.5^ for each parameter), likely attributable to errors in the estimation of the sampling parameters.

## 5 Discussion

We have introduced and implemented a stochastic model of *intrinsic* transcriptional noise that accounts for sequencing artifacts or *technical* noise. This model addresses an apparent overrepresentation of long unspliced mRNA in a variety of scRNA-seq datasets, and we posit that this bias is unlikely to arise biologically: fitting a simple model of mRNA production, splicing, and degradation produces parameter trends that render the fits suspect. Instead, we propose a model motivated by the chemistry of the sequencing process: each mRNA can be captured and reverse transcribed multiple times, with the possibility of such false positives growing with the length of molecule and the number of poly(A) capture sites (Fig. S2).

We fit this model to a variety of datasets, and discover that the parameter values, and thus entire mRNA distributions, are consistent for sets of technical and biological replicates. Further, the parameter values themselves (Fig. S12) are concordant with previous reports. Average burst sizes in the technical noise model are in the range (10^0.5^, 10^1.5^) [27, 28], rather than (10^−0.5^, 10^0.5^) in the noise-free model (Fig. S5). Degradation rates *γ*/*k_i_* are in the range (10^0^,10^1^), roughly consistent with fluorescence-based genome-wide results [4]. Finally, the splicing rates *β*/*k_i_* are relatively slow and largely fall in the range (10^−0.5^, 10^0.5^), on the order of 100 min. This result suggests that *β* is best interpreted as the rate of an abstracted, multi-intron process, as a single intron takes minutes to tens of minutes to splice [16, 17, 19]. We discuss potential refinements of this model in Section 5.2.

While the model performs well with whole-cell scRNA-seq data, it yields poor fits and inconsistent parameter values when applied to a nuclear mRNA datasets (Fig. S11). This phenomenon can be interpreted mechanistically as a reduction in degradation. Instead, the efflux of spliced mRNA is purely due to *export* from the nucleus to the cytoplasm. Therefore, we suggest that the dramatic increase in inferred degradation rates appears to reflect a rapid efflux process, whereas the high rate of reported model rejection may be explained by misspecification. For example, this efflux may be non-Markovian, and occur after a deterministic, rather than stochastic, delay. Unfortunately, in spite of extensive recent analytical work [29–34], probabilistic solutions are only available for constitutive, not bursty systems [5], making this hypothesis challenging to test.

We propose that our approach has applications beyond technical batch effect regression: it offers a mechanistic approach to the identification of differentially regulated genes. Instead of testing differences in average expression [7], our model enables testing differences in *parameter values*. This conceptualization likely provides multiple advantages. Firstly, it increases statistical power, due to reliance on model-specific results rather than non-parametric limiting theorems. For example, a gene may be expressed at nearly identical average levels in two cell types, but have very different distributions; such an effect is easier to detect using full parametric distribution fits. Secondly, it yields greater interpretability, as all parameters explicitly model biophysical processes. For example, a difference in average expression may be directly attributed to modulation of burst size or a reaction frequency, as in previous work using fluorescence-based measurements [35]. We provide a sample comparison between mouse heart and brain tissue inference results in Fig. S13, which demonstrates weakly correlated, often discordant parameter fits. More sophisticated theoretical machinery must be developed to quantitatively compare different tissues or experimental conditions: the offsets seen in Fig. S13 suggest that the sampling parameter fits are suboptimal, and thus skew estimates of splicing and degradation parameters. Multiple solutions are available, ranging from *ad hoc* selection of sampling parameters that concord with the identity, to fully Bayesian estimation of the posterior distributions.

### 5.1 Previous work

Wang et al. [36] use a similar sampling model, with *X*|*n* ~ *Poisson*(*Cn*). Their description recognizes the concordance between Poisson and Bernoulli models at low sampling rates. However, the underlying motivation for the Poisson model is obscure, and not motivated by the possibility of multiple priming for a single transcript. The authors make reference to Kim et al. [37], which models the number of cDNA molecules available for sequencing using a Bernoulli model. Although the same manuscript posits that the number of observed transcripts is Poisson-distributed with respect to the cDNA count, no technical justification is provided, nor is any mechanistic hypothesis advanced to explain the apparent emergence of new artifactual UMIs. The study only considers exonic reads and models them as direct gene products; as a consequence, it cannot represent buffering by splicing, which has been implicated as a significant mode of copy number control [38]. More recent work by Sarkar and Stephens [39] likewise posits a Poisson model, motivated by an approximation of the multivariate hypergeometric distribution [40] previously used to model relative abundances in the context of non-UMI technologies [41], rather than the details of sequencing. In contrast, we develop a concrete physiological and technical multimodal model with reference to the chemistry of the underlying sequencing process.

The aforementioned studies generally omit detailed discussion of the intrinsic biological noise. Several reports do consider technical and biological noise in scRNA-seq datasets, although with limited discussion of the origin of technical noise. The D^3^E package fits the one-species telegraph model, with analysis of sensitivity to dropout events [42]. However, the full drop-out model was not solved or fit and the inference procedure did not attempt to identify the technical noise parameter. Furthermore, the parametric form (dependent on the *biological* average expression) describes *cellular* drop-out events [43], without clear physical motivation: it is not obvious why the same gel bead may have UMIs corresponding to one gene but a total loss of the other. Further, these models do not have a self-consistent multi-gene interpretation: even if the dropout events are to be treated as cell-wide failure in cDNA library construction, the model does not imply that the *same* cells fail for different genes. Although this “zero-inflated” model has been conventional in scRNA-seq analysis [10,43,44], it has undergone substantial criticism in recent years [40,45], and we suggest the theoretical issues outweigh the appeal of tractability.

Overall, our modeling approach concords with some of the recommendations made in a recent study by Kim et al. [46]: normalization, transformation, and imputation are useful tools for data exploration but they may severely distort the underlying data. As we discuss in Section 5.2, some cell cluster analysis is necessary when homogeneous cell populations are present. However, we argue that the Poisson model they suggest for a homogeneous cell population is not consistent with findings from orthogonal methods, such as the pervasive transcriptional bursting observed by fluorescence measurements [4,27,35,47]. The study’s focus on zero fractions as a test statistic for model identification is likewise problematic: despite apparent agreement between empirical and Poisson zero fractions, the actual RNA distributions are near-universally overdispersed (Fig. 1a of [46]). Therefore, we again emphasize the general need for mechanistic models consistent with orthogonal data and full distributions. However, it is plausible that the approach of Kim et al. may be appropriate for certain transcripts present at high abundance and serving as cell type markers. We report the zero fractions for unspliced species in Section S1.3.

### 5.2 Limiting assumptions and future directions

While the noise modeling provided by our model adds realism over noise-free CME models, it relies on several simplifications. We discuss these at length, describe the physical basis of the assumptions, and propose corrections which may be implemented to represent more domains of behavior in extensions of this work.

Most prominently, the entire population of cells is presumed *homogeneous* and *ergodic*.

The homogeneity assumption implies that there is a *single* population of cells, which are interchangeable. This means the model is unable to directly represent heterogeneous samples with multiple distinct cell types. In practice, this simplification reflects the standard assumption that a relatively small set of marker genes is differentially regulated between different subpopulations or cell types. We use the chi-squared test with the Bonferroni correction to test agreement with the model fit. Genes that show, e.g., substantial bimodality, are excluded from the analysis.

Several approaches can be used to describe cell heterogeneity. To start, we can assume the cell population consists of a set of discrete, disjoint cell types. A computationally facile “top-down” approach first identifies cell types using standard clustering workflows, then separately fits the parameters for each cell cluster. In contrast, a fully stochastic, “bottom-up” approach might use the expectation-maximization algorithm to assign cell identities in a mixture model.

Ergodicity assumes that the cell populations are in equilibrium. This means the model is unable to directly represent samples from differentiation pathways. In practice, this simplification reflects the assumption that only a small subset of marker genes drive the differentiation process, and can be excluded using statistical testing; most genes are presumed to be ergodic.

Nevertheless, cell differentiation can be conceptualized using multiple probabilistic models. If we assume the cellular processes are slow relative to the mRNA dynamics, we can model the trajectory by a set of discrete cell types and fit them under the assumption of local ergodic equilibrium. If orthogonal information about the direction of the cell type trajectory is available, this model allows the investigation of parameter evolution throughout a differentiation process. However, the assumption of local equilibrium makes it impossible to *fit* the trajectory. On the other hand, it is possible to model the cell types as deterministic or stochastic functions governing the parameter values, and fit data to the resulting non-ergodic *occupation measures*.

We do not normalize by cell sizes at any step in the process, as normalization makes the data incompatible with the CME framework. Although this choice stands in stark contrast to standard scRNA-seq workflows [7], it is rigorously motivated by the kinetics of the process. The production of unspliced mRNA is presumed to occur at a single gene locus in a zeroth-order reaction with no spatial dependence. The splicing reaction is first-order, and the binding of the spliceosome is presumed to be non-limiting, which is supported by the reaction’s localization within the nucleus. Finally, the degradation reaction is likewise first-order; the binding of RNase is assumed non-limiting, i.e., the concentration of RNase is modeled as uniform throughout the cell. More complex models that incorporate second-order reactions would necessitate accounting for the cell volume to represent the probability of two molecules encountering each other.

We do not account for cell sizes in the model of the sequencing process. This description is motivated by simplicity, but we can make it rigorous based on chemically plausible assumptions. A crucial assumption is required for cell size independence during the library construction process: the process must be homogeneous.

Homogeneity in sequencing simply means that the reaction rates are constant across all cells, as well as over the duration of the reaction. All experimental steps after encapsulation in a gel bead take place in a cell-free medium. Assuming sufficient quality control in bead, droplet, and reaction mixture preparation, each cell’s reagent concentration is identical, at least at first.

Whether it remains this way depends on the molecular concentrations and process kinetics. For example, if the amount of reagents and primers is much greater than the number of mRNA, even the hypothetical sequencing of every molecule in a cell would not substantially deplete the reagents. The number of sequencing primers on a bead tends to exceed the number of mRNA. A typical human cell has 10^5^ − 10^6^ mRNA [48–50], whereas the number of *unique* UMIs in 10X v3 chemistry is usually above 10^6^ [12]. Although the distribution of primer counts on the proprietary 10X beads does not appear to be characterized in the literature, it is well-known that inter-gene UMI collisions (identification of distinct molecules with identical UMIs in a given cell) frequently occur in 10X datasets [12], which implies at least a several-fold excess of primers. The enzymes are catalytic, so their concentration is unlikely to vary over the course of the reaction. Finally, under the assumption that reverse transcription reagents are not depleted, cell size bias is implausible. All steps after library construction take place in combined media, rendering the cell size irrelevant altogether.

We note that it is possible to construct models more sophisticated than the homogeneous pure birth process to describe library construction. For example, we may describe the gradual denaturation of enzymes by a decaying sampling rate. This turns out to make no difference to the mathematical formulation of the process: the distribution of cDNA produced by an inhomogeneous arrival process is still Poisson [51]. Quantitatively, this means that a cDNA synthesis reaction with duration *t* yields *M_z_*|*N_z_* ~ *Poiss*(*N_z_*Λ_*z*_), with 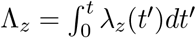 for some time-dependent rate λ_*z*_. Therefore, we can relax the assumptions to suppose such an *effective* sampling rate Λ_*z*_ is identical for all cells. We anticipate that multinomial models can provide a useful orthogonal description [40], but may be challenging to analyze jointly with Markov models of transcription.

Of course, cell sizes do vary in a non-stochastic way, and there is evidence that transcriptional parameters are in part regulated based on cell size [52]. More complex models may explicitly incorporate this dependence; yet again, there are several approaches to account for coordinated expression levels across multiple genes. A “top-down” model may use the total number of mRNA in a cell as a proxy for cell size, as is standard practice [7, 36], and represent the burst parameters as a mixture model. On the other hand, a “bottom-up” model may explicitly describe burst size synchronization between genes [5]. However, such models are rather complex to formulate and solve, so we omit their discussion, and reiterate that results presented here suggest fair agreement with the model and no further unexplained sources of overdispersion.

We use statistical testing to exclude genes that do not fit the bursty model. However, it is possible to use information criteria to identify other transcriptional models. For example, it is straightforward to model constitutive production [53], potentially with parametrized extrinsic noise [54, 55]. Sampling of these models yields Poisson-negative binomial (or Poisson-Poisson) mixtures easily tractable using the generating function method [56]. Furthermore, analogous solutions are available for the Poisson-Beta telegraph model [57]. However, empirically, the Poisson and Poisson-Beta distributions are rarely observed in scRNA-seq datasets [10], and bursty production is supported by fluorescence studies [4, 27]. These considerations, and the considerable computational expense of fitting multiple models for each gene, lead us to focus on the current model.

The binary model classifying each molecule as either spliced or unspliced is computationally tractable, but unsatisfying: it is only mechanistically justified in the case where only one intermediate transcript and one protein-coding transcript are prevalent. However, differential isoform expression is well-known to be prominent [25,58], with significant biological impact [59,60]. In principle, a more sophisticated workflow may identify individual intermediate isoforms and fit a full splicing graph. At this time, two protocols are available for isoform identification: Smart-seq3 [61] and FLT-seq [58], which generate full-length libraries with cell barcodes and UMIs. However, these methods can only capture relatively small numbers of cells, rather insufficient for full splicing graph and parameter estimation. Further, their annotations are not directly applicable to short-read data: short reads can identify *classes* of compatible transcripts rather than full transcripts. Therefore, the problem of fitting full isoform dynamics is statistically challenging. Parenthetically, we note that both methods use poly(A) capture, and are expected to yield capture biases similar to the short-read protocol.

The deterministic dependence on length is a simple model used as a proxy for the number of poly(A) tracts. With the growing availability of isoform-specific data, it is possible to build a more detailed thermodynamic description that explicitly models the rate of molecule capture as a function of the poly(A) content. The current implementation is partially modular to such descriptions, and permits using thermodynamic binding propensities instead of gene lengths.

Finally, we focus on maximum likelihood parameter estimates. Due to the high dimensionality of the problem, and relatively high computational cost of evaluating the relevant distributions, it is impractical to compute true confidence sets in the full search space. We implement a routine that approximates *conditional* confidence intervals for maximum likelihood {*b*, *β*, *γ*} vectors (at a given {*C_u_*, λ_*s*_}) by evaluating the Hessian of the KL divergence. With sufficient computing power, it is feasible to use standard Markov chain Monte Carlo schema to sample full posteriors.

## 6 Data and code availability

https://github.com/pachterlab/GP_2021_3 contains a Python notebook that can be used to reproduce the figures, as well as a sample notebook that applies the computational pipeline to a 10X PBMC dataset. The same repository contains all scripts used to make references, download datasets, quantify transcripts, and process the resulting *loom* files through the inference pipeline. The raw *loom* files and all search results are deposited in the CaltechDATA repository [62, 63].

## 7 Acknowledgments

G.G. and L.P. are partially funded by NIH U19MH114830. The DNA and RNA illustrations used in Figures 1 and 2 are derived from the DNA Twemoji by Twitter, Inc., used under CC-BY 4.0.

## Supplementary Material

### S1 Stochastic model of gene expression and sequencing

#### S1.1 Model derivation

The gene-specific model of transcriptional physiology assumes the existence of a single gene locus with stochastically regulated expression. Bursts of gene transcription arrive at exponentially distributed intervals with rate *k_i_*. This model can be derived from the more general two-state telegraph model of non-leaky gene expression at a locus:

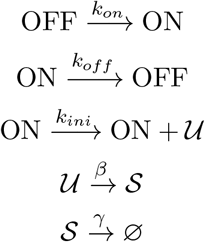

In the limit of *k_off_* → ∞, *k_ini_* → ∞, with *k_ini_*/*k_off_* → *b* ∈ (0, ∞), the three-parameter telegraph model reduces to the two-parameter burst model, yielding geometrically-distributed bursts with expectation *b* at each transcription event [21]. The burst frequency is simply *k_i_* ← *k_on_*. The unspliced mRNA 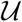 is converted to spliced mRNA 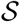 after an exponentially distributed interval with rate *β* the spliced mRNA is degraded after an exponentially distributed interval with rate *γ*. Since the single-cell RNA sequencing data are intrinsically atemporal, i.e. no natural experimental timescale exists to determine absolute rates without explicit experimental design, such as 4-thiouridine labeling of newly synthesized mRNA [8,64], the rates *β* and *γ* are only considered in units of *k_i_*, the burst frequency. Equivalently, *k_i_* is set to 1.

An expression for the system’s probability generating function (PGF) *G*(*g_u_*, *g_s_*, *t*) is readily available [6]. By evaluating *g_u_* and *g_s_* around the unit circle and performing an inverse discrete Fourier transform, it is straightforward to evaluate the time-dependent probability distribution of unspliced and spliced mRNA copy numbers (*P*(*n_u_*, *n_s_*, *t*). Due to the aforementioned atemporality of the scRNA-seq data, we only consider the steady-state distribution as *t* → ∞, henceforth referred to as *P*(*n_u_*, *n_s_*). The Markov chain representing the traversal is irreducible and aperiodic, so a unique stationary distribution is guaranteed to exist.

The mRNA population is presumed distributed according to *P*(*n_u_*, *n_s_*). We model the cDNA library construction from species *z* as a pure birth Poisson process with rate *D_z_*. This choice is motivated by the chemistry of the process [20]. We assume that the fixation of the cell medium stops all transcription, splicing, and degradation. Furthermore, diverging from previous descriptions [12], we model the process of the cellular mRNA being stripped off the newly synthesized cDNA by the template-switched second strand: the mRNA is not sequestered, and remains free to participate in further reactions. This makes the process catalytic, and suggests that the appropriate functional form for the distribution of cDNA produced from a single mRNA is a Poisson, rather than Bernoulli, distribution. Specifically, we posit that the rates of capture are usually sufficiently small that the Poisson probability of producing two or more cDNA is low; the correction is only necessary when the Poisson distribution rate is sufficiently high, such as when intronic regions have a large number of poly(A) priming sites.

Independent Poisson sampling yields cDNA copy numbers distributed per *Poisson*(*D_z_*) for each mRNA molecule, where *D_z_* is an experiment-specific sampling rate for species *z*. From standard stability properties, the distribution of the number of cDNA generated from *n_z_* mRNA is governed by *M_z_*|*n_z_* ~ *Poisson*(*D_z_n_z_*). By the law of total probability and the assumption of independent sampling processes, the full joint distribution is:

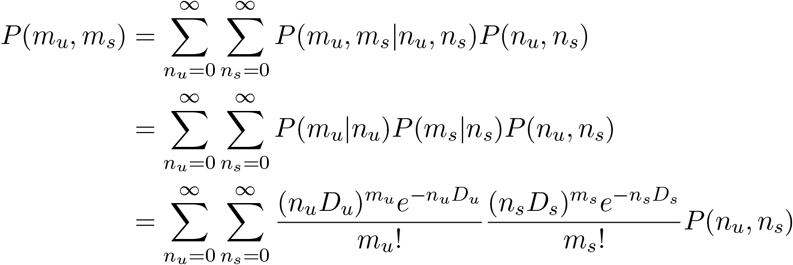

By the definition of the PGF of *P*(*m_u_*, *m_s_*):

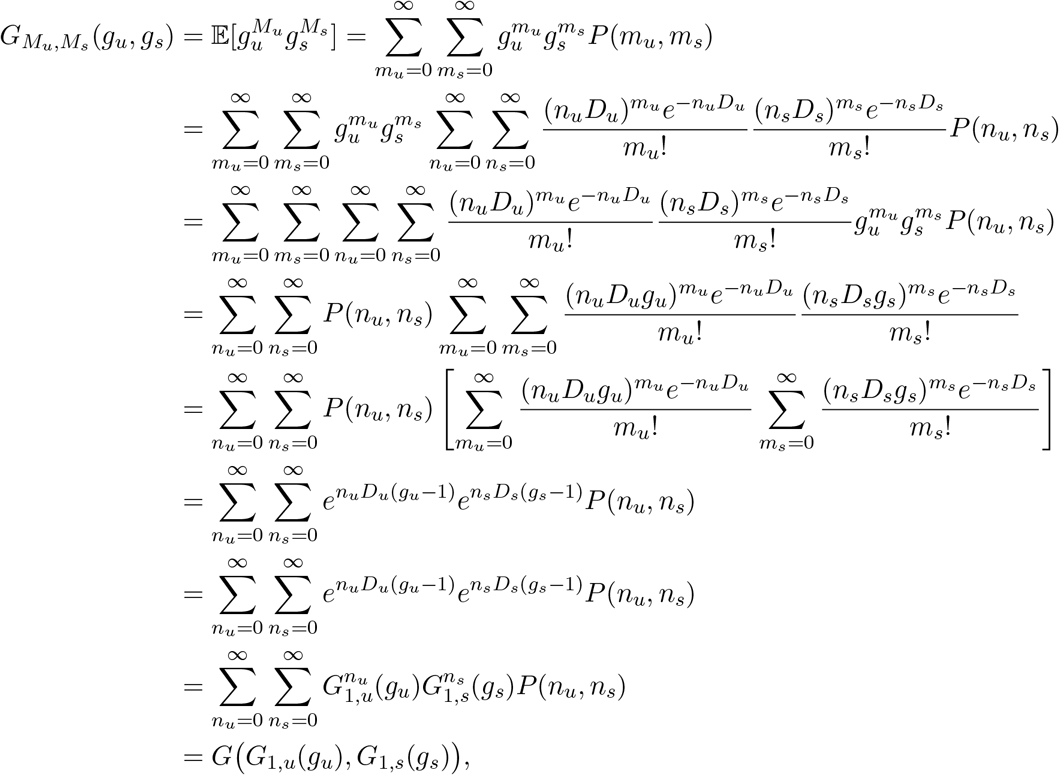

where *G*_1,*z*_(*g_z_*) is the PGF of the Poisson distribution with rate *D_z_* and the interchange of summation operators holds due to Fubini’s theorem. The final result accords with standard PGF identities.

After library construction, the cDNA molecules undergo amplification by polymerase chain reaction (PCR), fragmentation into short reads, adapter ligation, DNA sequencing, and identification from read data. The specific stochastic processes that govern these reactions are challenging to consider in detail. However, in contrast to the cDNA library construction, there appears to be no compelling mechanistic reason to presuppose any of the reactions can create novel false UMIs: previous studies have shown UMI collapse is effectively eliminates the transcription errors that arise in this process [12]. Therefore, as a first-order model, we treat these steps as a sequence of depletions or filters, amounting to *N_f_* steps of Bernoulli sampling applied to each molecule, with respective parameters 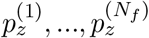. Trivially, this implies that the final per-molecule distribution is again Bernoulli, with a product probability 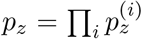. From standard identities, the number of observed UMIs, given the existence of *m_z_* cDNA, follows *X_z_*|*m_z_* ~ *Bin*(*m_z_*,*p_z_*).

From standard properties:

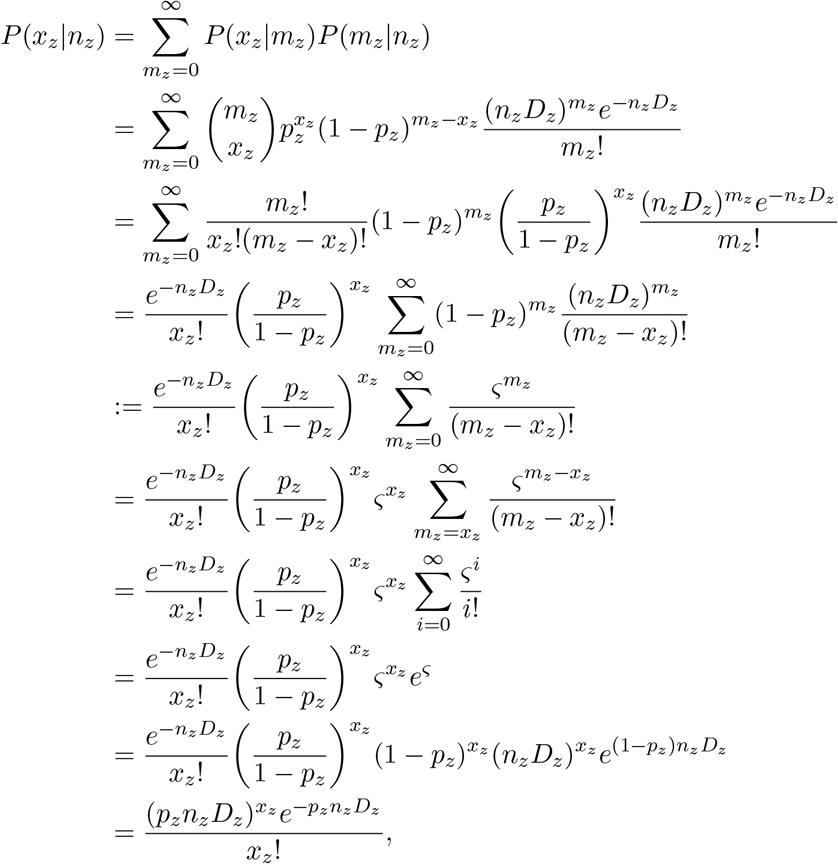

a Poisson distribution with a rate rescaled by the sequencing probability.

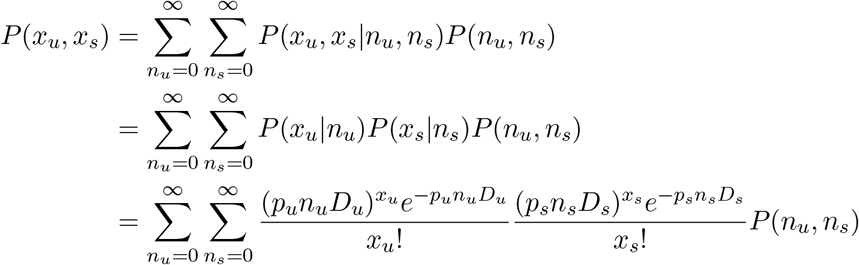

By the definition of the PGF associated with *P*(*x_u_*,*x_s_*):

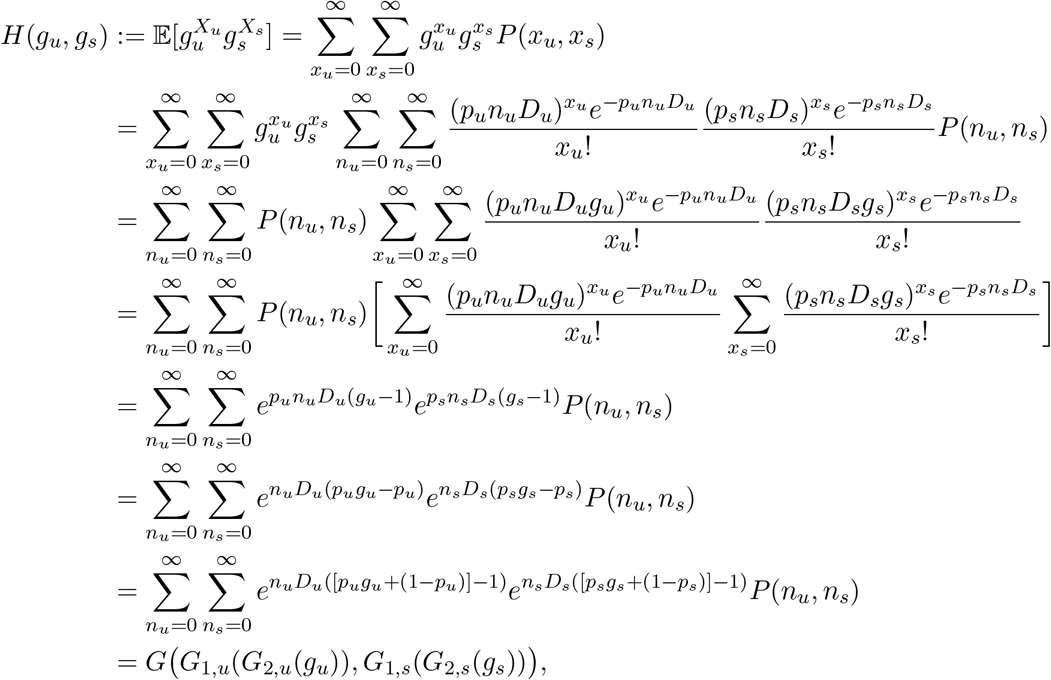

where *G*_2,*z*_(*g_z_*) is the PGF of the Bernoulli distribution corresponding to species *z* and the interchange of summation operators holds due to Fubini’s theorem. Again, this accords with standard properties of PGFs.

Most significantly to the formulation of the problem, the functional form of the results reinforces the fact that Bernoulli resampling of the distribution makes the individual parameters in the pairs *D_u_*, *p_u_* and *D_s_*, *p_s_* impossible to distinguish. Therefore, we define the effective capture rates λ_*u*_ ≔ *D_u_p_u_* and λ_s_ ≔ *D_s_p_s_*, such that each molecule of species *z* yields a Poisson distribution of observed UMIs with sampling rate λ_*z*_.

#### S1.2 Model moments

##### S1.2.1 Marginal moments

The moments of the marginals are easily acquired from standard descriptions of Poisson mixtures [65, 66]; here, we derive them explicitly using standard properties of generating functions. Given *H*(*g_u_*, *g_s_*) = *G*(*e*^λ*u*(*g_u_* − 1)^, *e*^λ_*s*_(*g_s_* − 1)^), the moments are found by taking derivatives at *g_u_*, *g_s_* = 1:

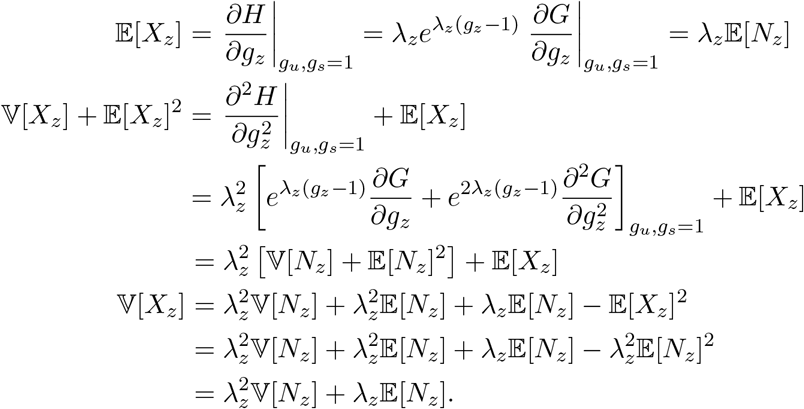

Usefully, this formula is independent of the specific form of *G*, and can be applied to any model of transcription and processing. In our case, per previous results [6]:

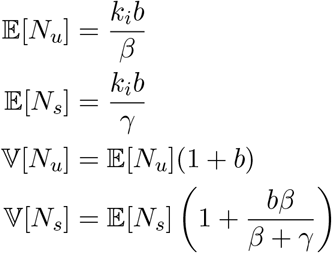

This yields the following analytical expressions for the moments of observables *X_z_* (defining 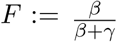):

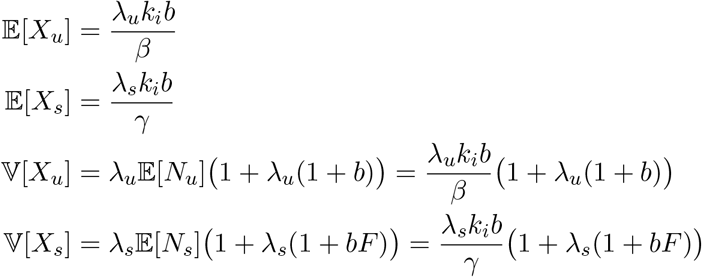

##### S1.2.2 Cross-moments

Analogously to the previous section:

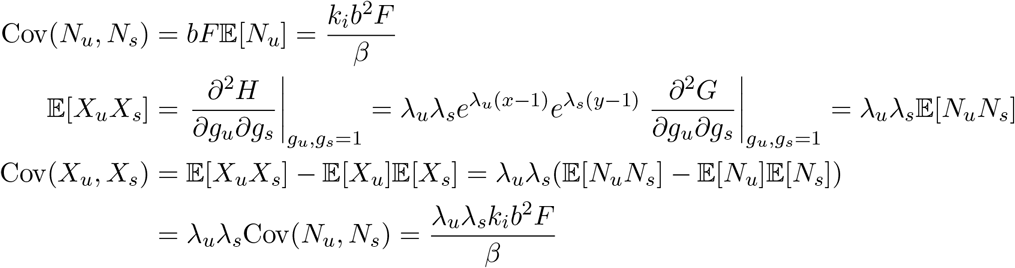

The Pearson correlation can be computed accordingly. First, we find the noise-free correlation *ρ*\*:

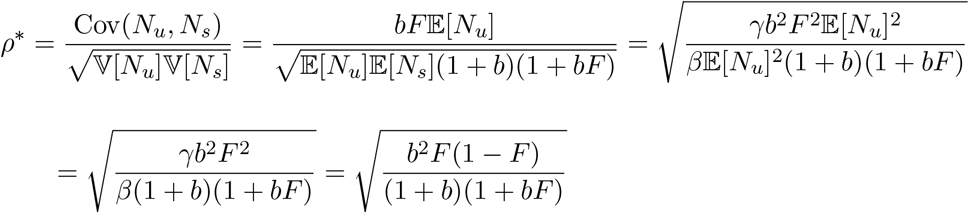

Then, we compute the correlation *ρ* of the sampled system:

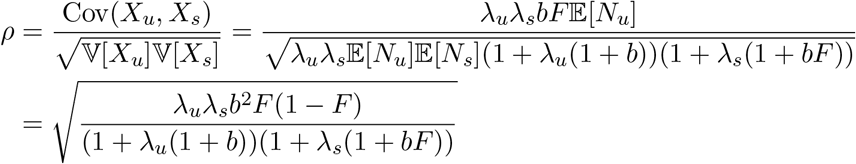

We can compare these quantities:

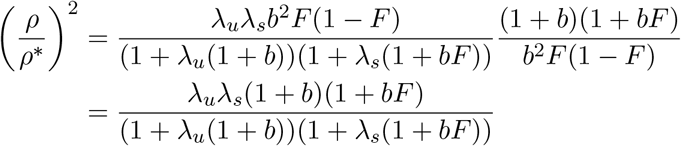

Defining ζ ≔ 1 + *b* > 1 and *η* ≔ 1 + *bF* > 1 yields:

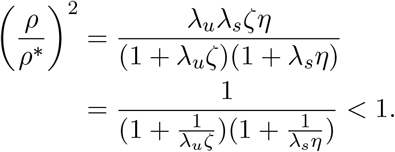

As expected, sampling strictly reduces the correlation with respect to the technical noise-free system, although the ratio of the correlation coefficients tends toward 1 as λ_*u*_ζ and λ*_s_η* tend toward infinity.

##### S1.2.3 Method of moments parameter estimates

We set *k_i_* to 1 with no loss of generality at steady state. Treating the noise-free model, we can easily compute method of moments estimates for the three physiological parameters using the analytical results in Table 4:

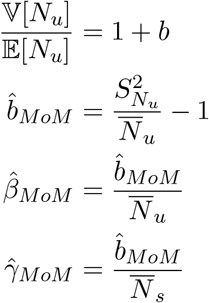

Given λ*_u_* and λ*_s_*, we can compute the conditional method of moments estimates for the three physiological parameters in the Poisson technical noise model:

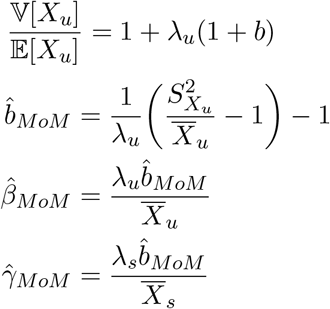

#### S1.3 Zero fractions

Due to the sparsity of molecular datasets, the zero fraction *P*_0_ is a popular data summary [21,46,67]. In general, we suggest using fully parametric fits, but we report the *P*_0_ for the considered models. The zero fraction is the value of the distribution’s PGF at *g_z_* = 0. Therefore, for a Poisson distribution with mean λ, 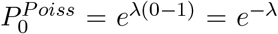. The same relation can be applied to all other PGFs. We omit the discussion of spliced species, as they do not have analytically tractable solutions, but we do note that computing them requires only a single integral

First, we consider the original bursty system.

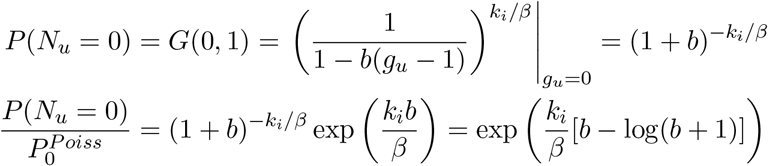

Since *b* > 0, the argument [*b* – log(*b* + 1)] is strictly positive. This implies that the zero fraction of the bursty system is strictly higher than that of the moment-matched Poisson system.

In the case of the sampled system:

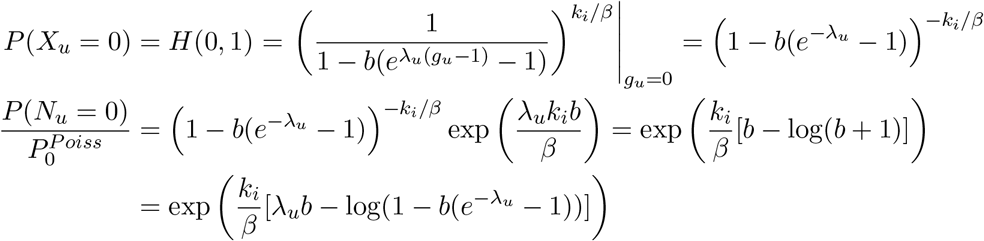

Since *b*, λ_*u*_ > 0, the argument [λ*_u_b*−log(1 –*b*(*e*^−λ_*u*_^ − 1))] is strictly positive. Again, this implies that the zero fraction of the sampled bursty system is strictly higher than that of the corresponding Poisson system. These results provides a route to computing the false positive rate, if model selection is undertaken using the zero fraction [46].

#### S1.4 Sampling parameters

At this point, the specific functional forms of λ*_u_* and λ*_s_* are left to discretion; each could be gene-specific or universal across the transcriptome. Due to the behavior of the summary statistics of experimental data, we hypothesize that the spliced mRNA capture rate λ_*s*_ is constant for all genes, whereas the unspliced mRNA capture rate λ_*u*_ has a linear dependence on the gene length, such that λ*_u_* = *C_u_L*.

The identification of intronic sequences is presumably [8] enabled by the off-target capture of intronic poly(A) sequences. This effect has been known as a source of significant systematic bias since at least 2002 [22]. A study by Nam et al. found that a poly(A) stretch of length eight – with up to two internal mismatches – was sufficient to initiate priming; increasing the stretch length substantially increased priming efficiency. We do not take the specific poly(A) content of each gene into account, but note that our computational implementation allows using the number of poly(A) stretches up to a specified length instead of *L*.

### S2 Simpler models produce implausible parameter trends

To motivate the need for the length-biased Poisson sampling model, we consider a series of simpler models: the aforementioned noise-free model, the Bernoulli sampling model, and the length-independent Poisson sampling model, and find that all lead to the counterintuitive length-dependent trends characterized in Section 2.1.

#### S2.1 No sampling

As described in Section 2, the current study is motivated by the incongruity between the genomewide parameter trends known from previous literature and inferred from typical scRNA-seq datasets. To compute the parameter estimates, we simply perform maximum likelihood estimation on the same data, initializing at the method of moments estimates reported in Section S1.2.3.

As shown in Fig. S5, the results are mutually concordant, and physiologically implausible, across a variety of high-quality scRNA-seq datasets.

#### S2.2 Bernoulli sampling

Given these results, we may reasonably suppose that a model with no technical noise is too simple, and posit that the library construction and sequencing processes can lose molecules. This pure-depletion Bernoulli sampling model has the following per-molecule generating function:

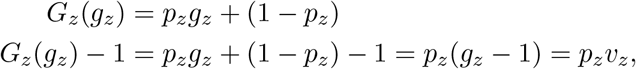

which implies that the overall PGF has the following functional form:

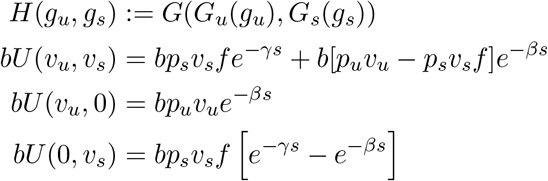

Clearly, binomial sampling of the marginal distributions returns the same functional form, with effective burst size *bp_z_*. However, the joint distribution only takes an identical form if *p_u_* = *p_s_*.

For simplicity, we only consider the unspliced marginal. From Fig. S5, it is clear that the resulting qualitative parameter trends under the length-independent Bernoulli model must be identical, up to rescaling of the burst size by *p_u_*.

#### S2.3 Length-independent Poisson sampling

Even if we use the chemical considerations to suppose that Poisson sampling is necessary, it is not immediately clear that a length-dependent model is required. Considering only the unspliced marginal again, and supposing that λ*_u_* ≪ 1, we yield:

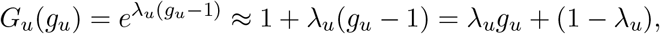

which is identical to the Bernoulli case, with *p_u_* ← λ*_u_*.

It is not *a priori* obvious that λ*_u_* ≪ 1 should be true. We fit a 1000-gene subset of the `pbmc_10k` dataset on a 22 × 23 grid, with log_10_λ*_u_*, log_10_λ*_s_* ∈ [−3.5, 1], and all other parameter bounds as in Table S3. The procedure discovered the sampling parameter optima (log_10_λ_*u*_, log_10_λ_*s*_) = (−0.93, −0.84) and the parameter trends shown in Fig. S14. These trends are essentially identical to the noise-free model (Fig. S5), up to translation due to scaling. Further, the low values of the sampling parameters indicate that the sampling distribution is in a Bernoulli-like regime, and support the qualitative results in Section S2.2.

### S3 Optimization and analysis

#### S3.1 Method of moments initialization

We initialize the maximum likelihood estimation algorithm at the method of moments (MoM) estimate. However, it is not immediately clear that a single search is sufficient for three-dimensional optimization over {*b*, *β*, *γ*}. There may be a risk of finding suboptimal local minima. To validate this choice, we tested whether the MoM initialization produces an improvement over random initialization, and whether twenty independent searches produce an improvement over one search. In the case of twenty searches with MoM, we initialized a single search at MoM and sampled all other starting points from a uniform distribution over the search space. We used these conditions to fit a 1000-gene subset of the `pbmc_10k` dataset on a 22 × 23 grid, with log_10_*C_u_* ∈ [−8.5, −3], log_10_λ*_s_* ∈ [−3.5, 1], and all other parameter bounds as in Table S3. To benchmark the standard setting (MoM, one search) against the three others, we plotted the optimal divergence of each gene at each {*C_u_*,λ_*s*_}.

As shown in Fig. S3, the random, 1-iteration condition often underperformed the standard, whereas the MoM, 20-iteration condition always outperformed it. As expected, the random, 20-iteration condition performed slightly worse than the MoM, 20-iteration condition.

From the comparisons, it is apparent that the MoM, 1-iteration search is outperformed by the 20-iteration searches when the divergence is relatively high. Therefore, we follow up and investigate whether the underperformance of the standard settings can actually impact the inferred sampling parameter optimum. We color the values near the best estimate of this optimum (the set of {*C_u_*,λ*_s_*} in the first quartile of total divergence in the MoM, 20-iteration search). In this region, the 1-iteration search performs as well as the 20-iteration search (orange points in Fig. S3). Therefore, underperformance is largely restricted to the sections of the parameter landscape far from the optimum, and does not substantially affect the optimization results.

The moment-based starting point provides a high-quality estimate, comparable to numerous, costly random initializations. Adding more starting points has very little marginal benefit. Therefore, we strictly use a single moment-based estimate to initialize likelihood optimization throughout this study. The number of starting points and their location (random or MoM) can be set by the user of the implementation.

#### S3.2 Statistical testing

The optimization procedure yields a set of best-fit parameters 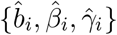 for each gene *i* and each {*C_u_*,λ_*s*_}, along with an accompanying gene-specific KL divergence or loss function *L_i_*(*C_u_*,λ_*s*_). Since each gene has an identical number of observations, we can ostensibly find a net *L*(*C_u_*, λ_*s*_) ≔ ∑_*i*_*L_i_*(*C_u_*, λ_*s*_) and estimate the optimal sampling parameters 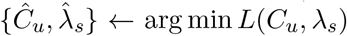. This approach is enhanced by testing and excluding genes that are poorly described by the model. Data may not fit well due to the simplifications discussed in Section 5.2, as well as convergence to suboptimal parameter vectors.

We use the chi-squared test with *p* = 0.05 and the Bonferroni correction to reject genes that do not appear to be accurately described by the model. Further, we reject fits that are too close to the biophysical parameter constraints (within 0.01 of the bounds), as they may represent degenerate cases or local optima.

However, rejecting a sufficiently large subset of genes may shift the global sampling parameter optimum. We do not account for this possibility in the analyses presented here, as multiple potential corrections are available. However, we do implement and make available several procedures to test the stability of the optimum.

The first accepts an integer number, repeatedly chooses a subset of genes of that size, and reports the landscapes and optima in the subsampled search results. The second seeks a *self-consistent* optimum: beginning at the naïve estimate of the sampling parameter optimum, it uses the chi-squared test to reject a subset of the genes, then computes a *new* optimum based only on the retained genes. The process repeats and reports the average of the optima observed throughout the process. We did not find the self-consistent correction to significantly shift the location of the optimum in the considered datasets.

### S4 Biological count estimation

Throughout the study, we omit the explicit discussion of *N_z_*. This approach is non-standard: a breadth of literature uses statistical models precisely to “regress out” technical noise effects, in practice taking *x_z_* and transforming it to some 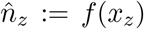, ostensibly corresponding to a (not necessarily integer) *in vivo* abundance. However, it is unclear how this abundance should be interpreted: 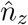 is an *ad hoc* point estimate, and assigning a single value to the physiological mRNA abundance obscures the loss of information in the sequencing process. In other words, every *x_z_* corresponds to an entire distribution of possible *n_z_*, and this distribution should be treated explicitly. Using a full mechanistic model obviates the need for describing it, because the relevant sums over the probability mass functions are taken during the generating function derivation (as in Section S1.1).

However, for the sake of completeness, we outline a procedure for estimating the physiological mRNA counts. Here, we consider the simplest case: we know the exact values of all parameters, and observe *x_z_* UMIs. Bayes’ theorem yields the distribution of biological counts:

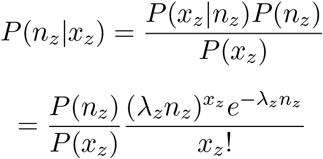

This expression is exact, and extending it to multiple species is trivial. However, no analytical solution is available. Furthermore, this form is not even computationally tractable, since no closed-form solutions for *P*(*x_z_*) and *P*(*n_z_*) are available: for sufficiently small λ*_z_*, the domain over which the PMFs must be evaluated grows considerably, making the first factor numerically unstable.

Nevertheless, there are some simple extensions which *are* analytically tractable. We can investigate the expectation of the distribution (replacing species z-specific notation with generic notation for convenience, but keeping in mind that the discussion concerns a single marginal species):

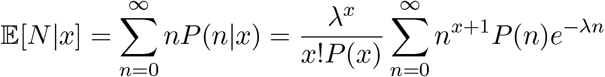

The rightmost term is recognizable as a derivative of the moment-generating function of *N*. Specifically, defining 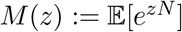, we yield:

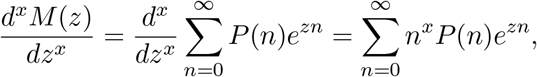

which immediately implies the identity:

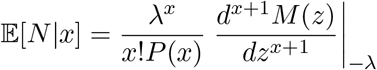

Analogously, it is possible to evaluate the PGF of *N*|x. Specifically,

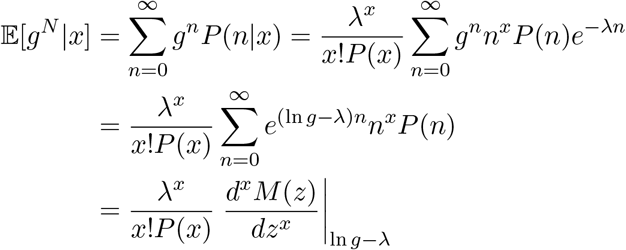

This form is relatively challenging to evaluate for large *x*, and requires the explicit computation of *P*(*x*). Nevertheless, we can treat some limiting cases explicitly, by defining a tractable functional form for *N*. First, we suppose *N* is distributed per *Poisson*(*θ*), i.e., the production is constitutive [53] or the molecule under study is a spliced mRNA species with very low *β* [6]. This yields a Neyman type A (Poisson-Poisson mixture) UMI distribution *X* [56]:

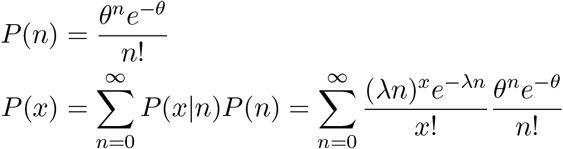

Applying Bayes’ theorem:

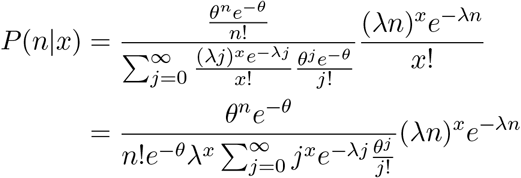

This expression can be computed explicitly. However, for the sake of qualitative investigation, we can consider the case *X* = 0, and compute the distribution of molecules implied by an observation of zero UMIs:

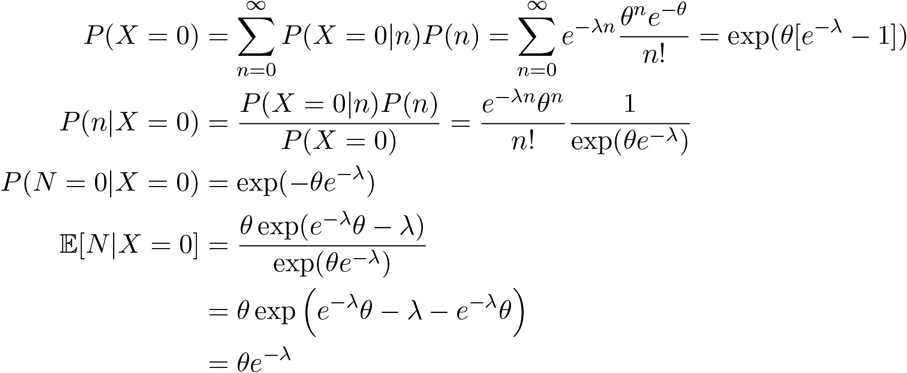

Therefore, if the sampling rate is high, we expect an observed zero to correspond to a true zero. On the other hand, if the sampling rate is low, the best estimate of the true abundance is simply an exponentially discounted average abundance. By Taylor expansion, this converges to a linearly discounted abundance as λ → 0.

As a second, more physiologically relevant illustration, we can consider the case where *N* ~ *NegBin*(*p,r*), i.e., the molecule under study is either unspliced, or spliced after a very brief delay (*β* ≫ 1). Supposing once again that zero molecules are observed:

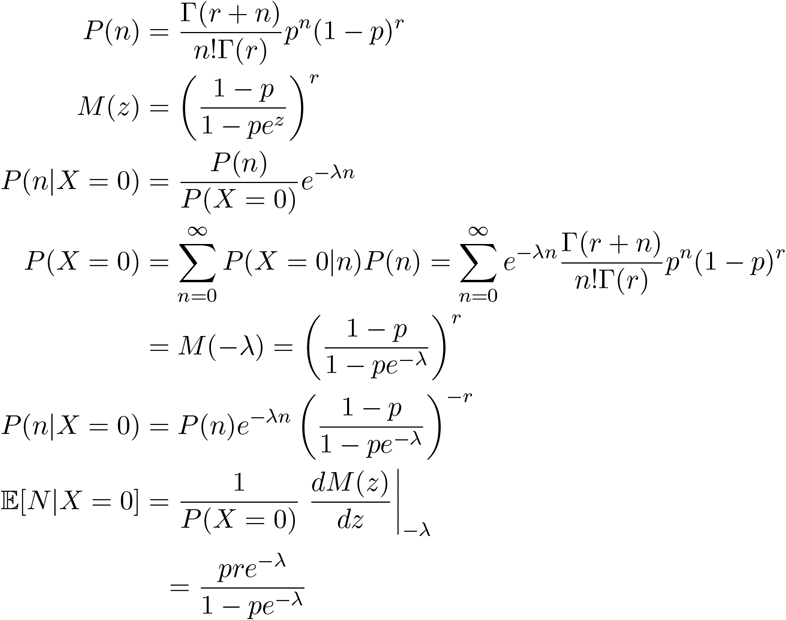

In terms of biological parameters 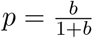 and *r* =*k_i_*/*β* [10], and recalling that the average biological expression 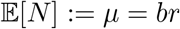:

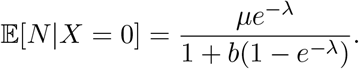

For high sampling rates, the expected physiological molecule number is vanishes: an observed zero is likely to be a real zero. As the burst size increases, the function reduces to *r*(*e*^λ^ − 1)^−1^. In the limit of low λ, this gives *r*/λ by Taylor expansion. Finally, for any *b* and low λ, we yield *μ* −*μ*(1 + *b*)λ, a scale-dependent correction to the mean.

In summary, it is possible to use the analytical models of transcription and sequencing, combined with Bayes’ theorem, to estimate underlying molecule counts. Some simple summaries are amenable to exact analysis; it is relatively straightforward to estimate the biological molecule counts implied by observing zero UMIs. Wherever analytical results are not available, it is possible (albeit computationally intensive) to approximate the solutions numerically.

However, this procedure is *not* model-agnostic “imputation” of missing values or zero observations: model and noise parameter estimates are required. Finally, we note that the theoretical analysis is performed under assumption of perfect information about the parameter values. However, extensions to standard Bayesian machinery are straightforward; in that case, the following formulation is appropriate:

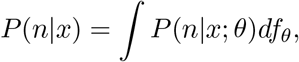

where *f_θ_* is the posterior distribution of inferred parameters, such as the approximate log-normal posterior discussed elsewhere in the report.

### S5 Supplementary tables

**Table S1:** Summary of datasets used in the analysis. The PBMC datasets originate from *H. sapiens*, all others originate from *M. musculus*.

| Dataset | Abbreviation | Cells | Genes Detected | Genes Kept | Genes Overlap | Genes Selected |
| --- | --- | --- | --- | --- | --- | --- |
| 10X 1k PBMC | pbmc_1k | 1200 | 36601 | 3087 | 2997 | 2500 |
| 10X 10k PBMC | pbmc_10k | 11756 | 36601 | 5813 |  |  |
| 10X 1k heart | heart_1k | 982 | 32285 | 6746 | 3607 | 2500 |
| 10X 10k heart | heart_10k | 7462 | 32285 | 8281 |  |  |
| 10X 1k neuron | neuron_1k | 1330 | 32285 | 4943 |  |  |
| 10X 10k neuron | neuron_10k | 11954 | 32285 | 5348 |  |  |
| 10X 5k brain | brain_5k | 5399 | 32285 | 5370 | 4775 | 3500 |
| 10X 5k brain nuclei | brain_5k_nuc | 5772 | 32285 | 5854 |  |  |
| Allen B01 | allen_B01 | 11504 | 32285 | 5894 | 5399 | 5000 |
| Allen C01 | allen_C01 | 12363 | 32285 | 6477 |  |  |
| Allen A08 | allen_A08 | 9974 | 32285 | 7367 |  |  |
| Allen B08 | allen_B08 | 10975 | 32285 | 7482 |  |  |

**Table S2:**
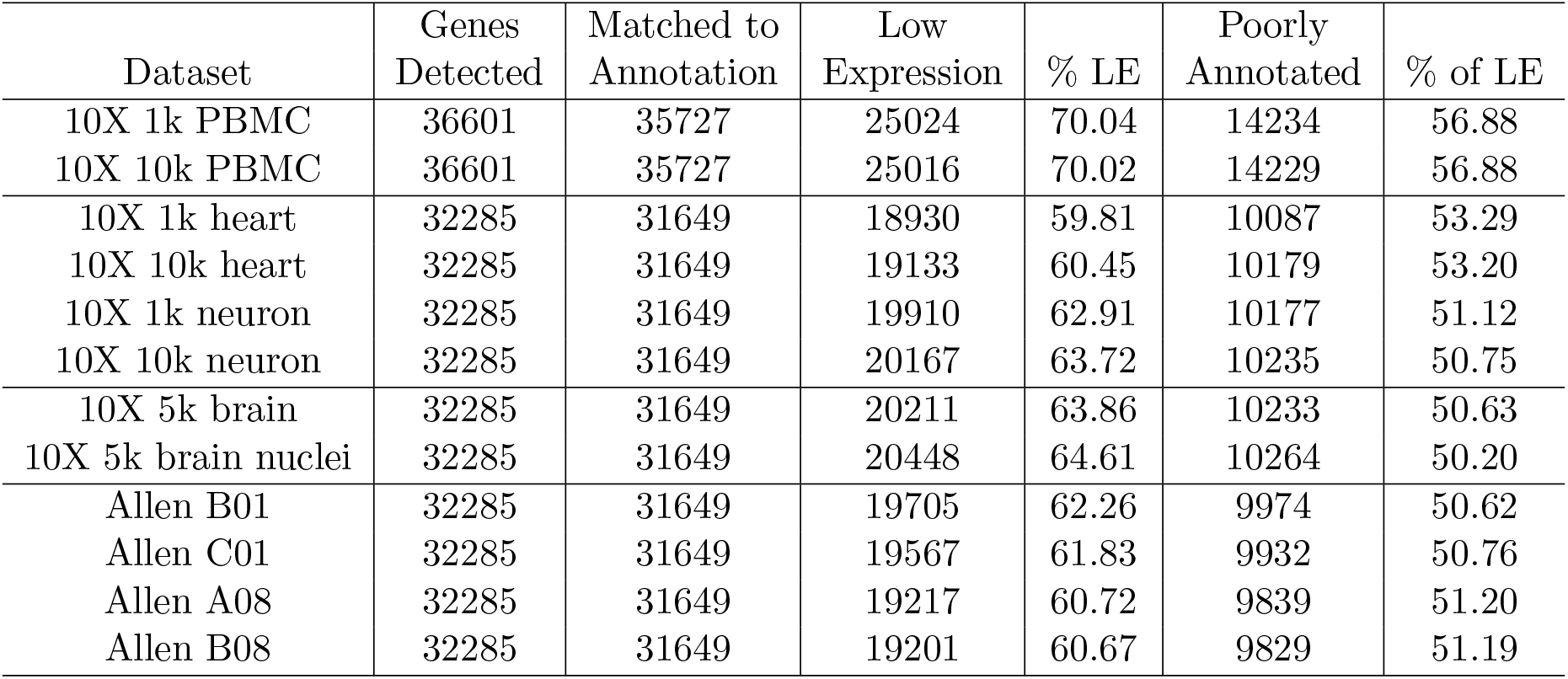
Analysis of the rejected low-expression gene cluster. The “Low Expression” cluster was assigned using K-means clustering. The “Poorly Annotated” genes were identified by parsing gene names to identify all genes known only as open reading frames with no functional characterization, or tentatively named using AC/AL/BC/Gm/LINC prefixes and Rik/AS/IT/ps pseudogene suffixes.

| Dataset | Genes Detected | Matched to Annotation | Low Expression | % LE | Poorly Annotated | % of LE |
| --- | --- | --- | --- | --- | --- | --- |
| 10X 1k PBMC | 36601 | 35727 | 25024 | 70.04 | 14234 | 56.88 |
| 10X 10k PBMC | 36601 | 35727 | 25016 | 70.02 | 14229 | 56.88 |
| 10X 1k heart | 32285 | 31649 | 18930 | 59.81 | 10087 | 53.29 |
| 10X 10k heart | 32285 | 31649 | 19133 | 60.45 | 10179 | 53.20 |
| 10X 1k neuron | 32285 | 31649 | 19910 | 62.91 | 10177 | 51.12 |
| 10X 10k neuron | 32285 | 31649 | 20167 | 63.72 | 10235 | 50.75 |
| 10X 5k brain | 32285 | 31649 | 20211 | 63.86 | 10233 | 50.63 |
| 10X 5k brain nuclei | 32285 | 31649 | 20448 | 64.61 | 10264 | 50.20 |
| Allen B01 | 32285 | 31649 | 19705 | 62.26 | 9974 | 50.62 |
| Allen C01 | 32285 | 31649 | 19567 | 61.83 | 9932 | 50.76 |
| Allen A08 | 32285 | 31649 | 19217 | 60.72 | 9839 | 51.20 |
| Allen B08 | 32285 | 31649 | 19201 | 60.67 | 9829 | 51.19 |

**Table S3:** Search parameter bounds. *Fit to `brain_nuc_5k` used [−0.5, 3.5] as the domain for log_10_ γ.

| Parameter | Lower bound ($\log_{10}$) | Upper bound ($\log_{10}$) |
| --- | --- | --- |
| $C_u$ | -8 | -5 |
| $\lambda_s$ | -2.5 | 0 |
| $b$ | -1 | 4.2 |
| $\beta$ | -1.8 | 2.5 |
| $\gamma^*$ | -1.8 | 2.5 |

**Table S4:**
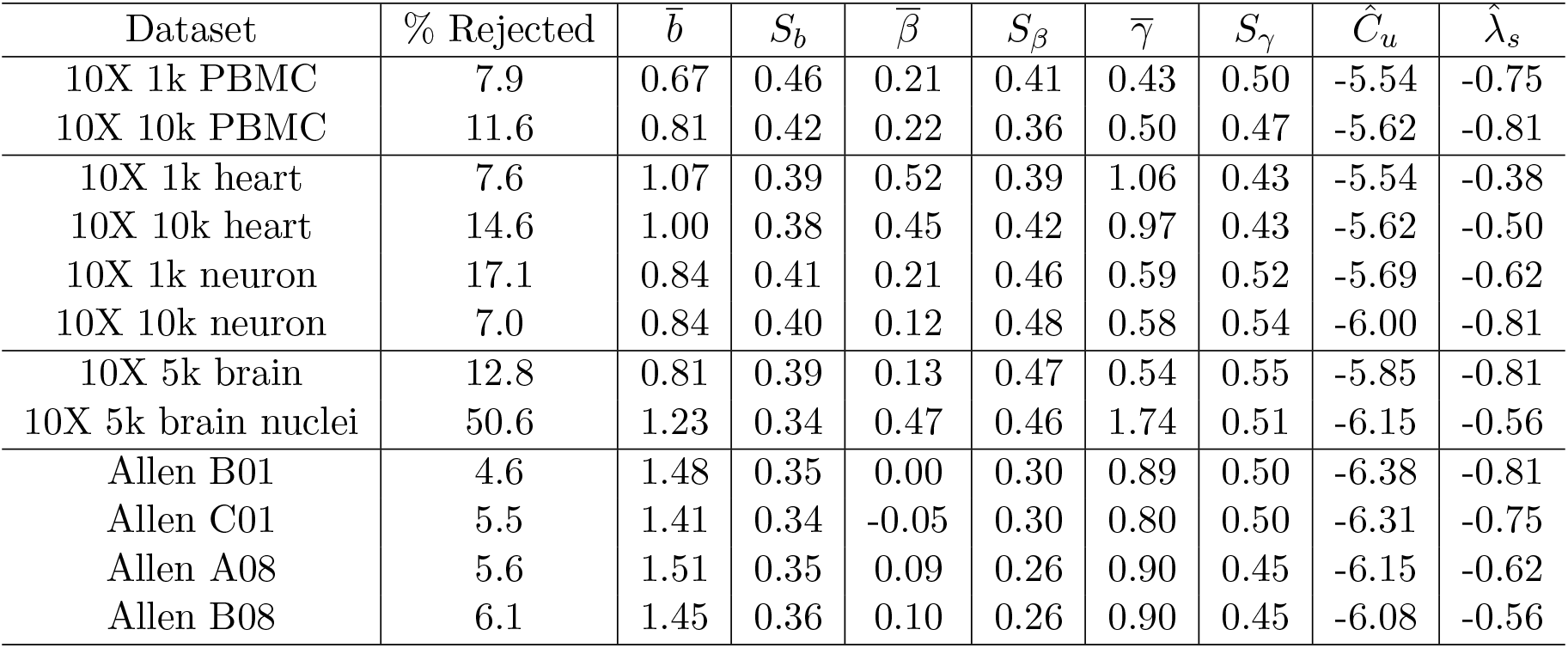
Statistical summaries of biophysical and technical parameter values computed by the inference procedure.

### S6 Supplementary figures

#### S6.1 Procedure and assumptions

**Figure S1:**
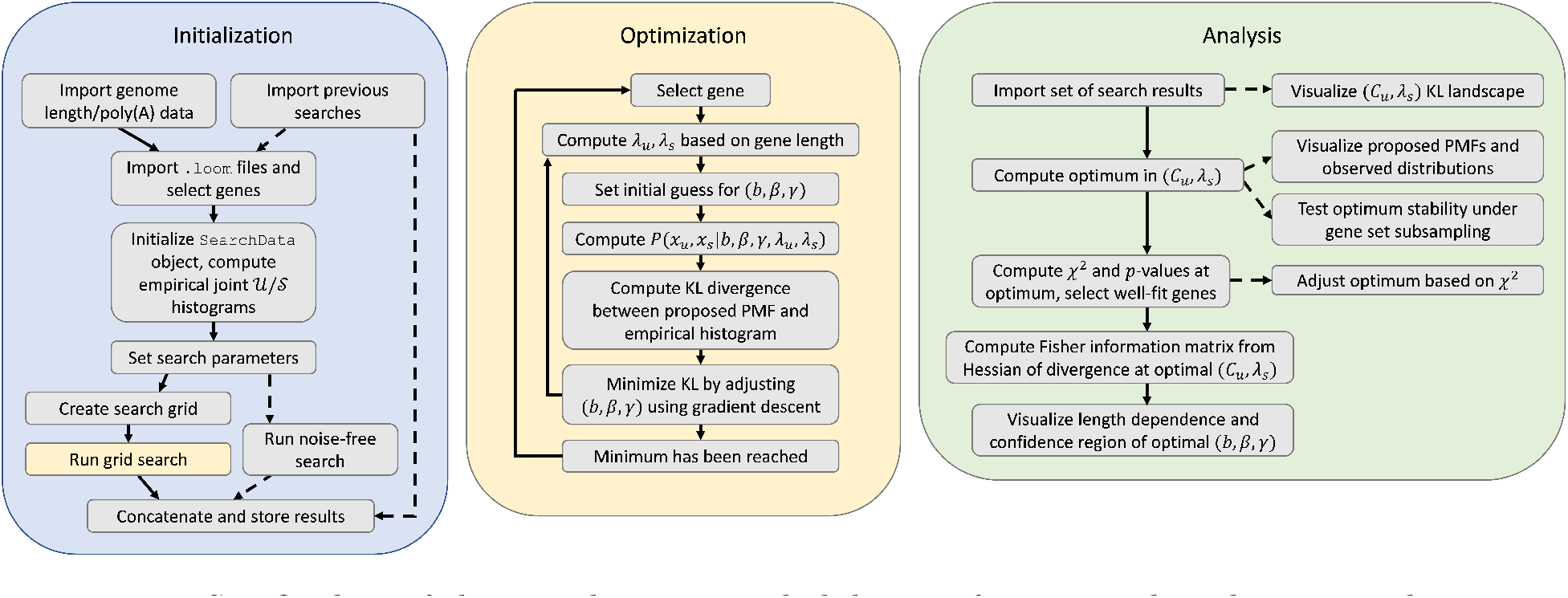
Outline of the initialization, probabilistic inference, and analysis procedure.

**Figure S2:**
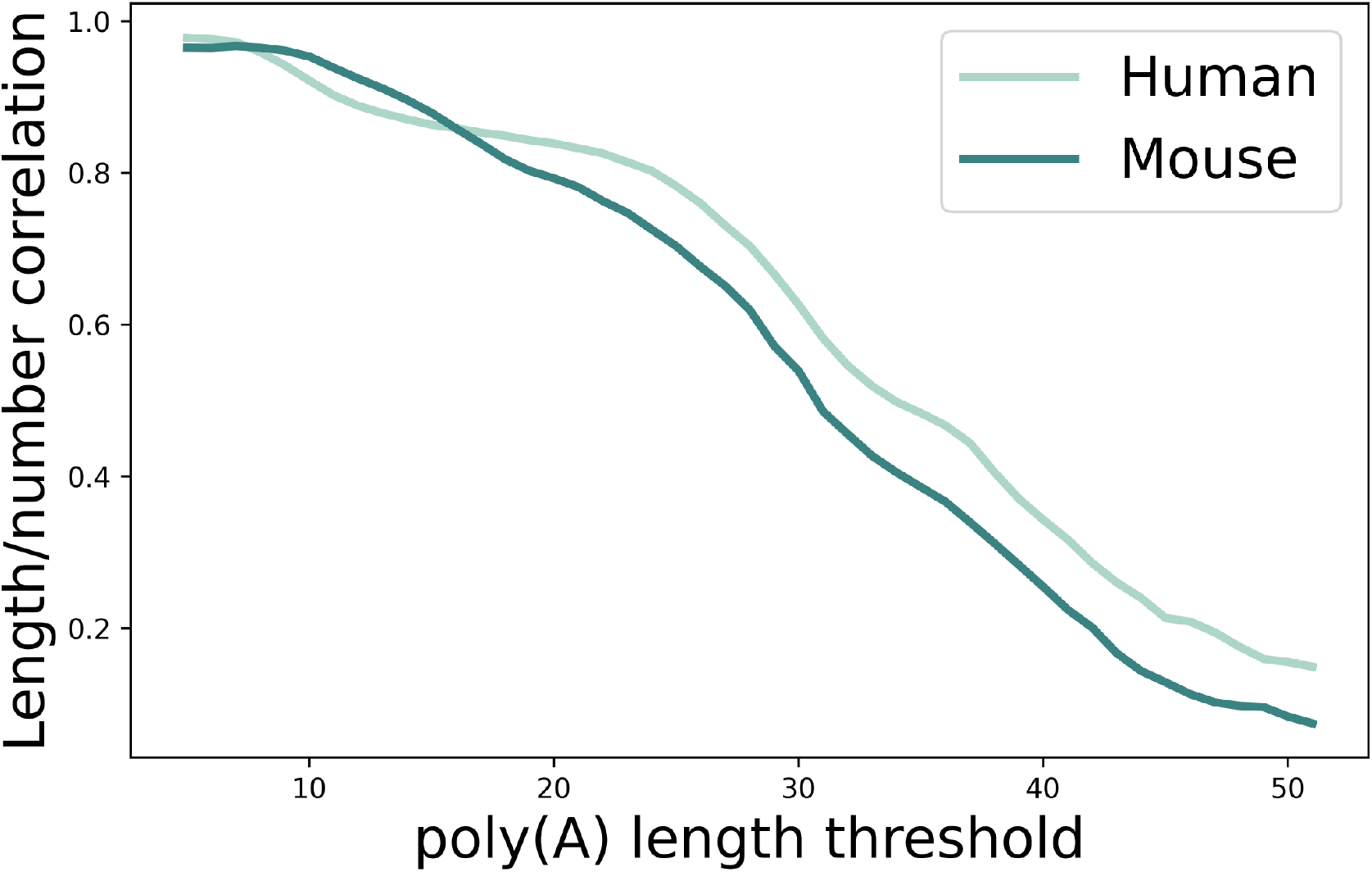
Correlation between the total gene length and the number of internal poly(A) stretches up to a set length. At the relatively low poly(A) stretch lengths necessary to initiate priming, the correlations are above 0.9.

**Figure S3:**
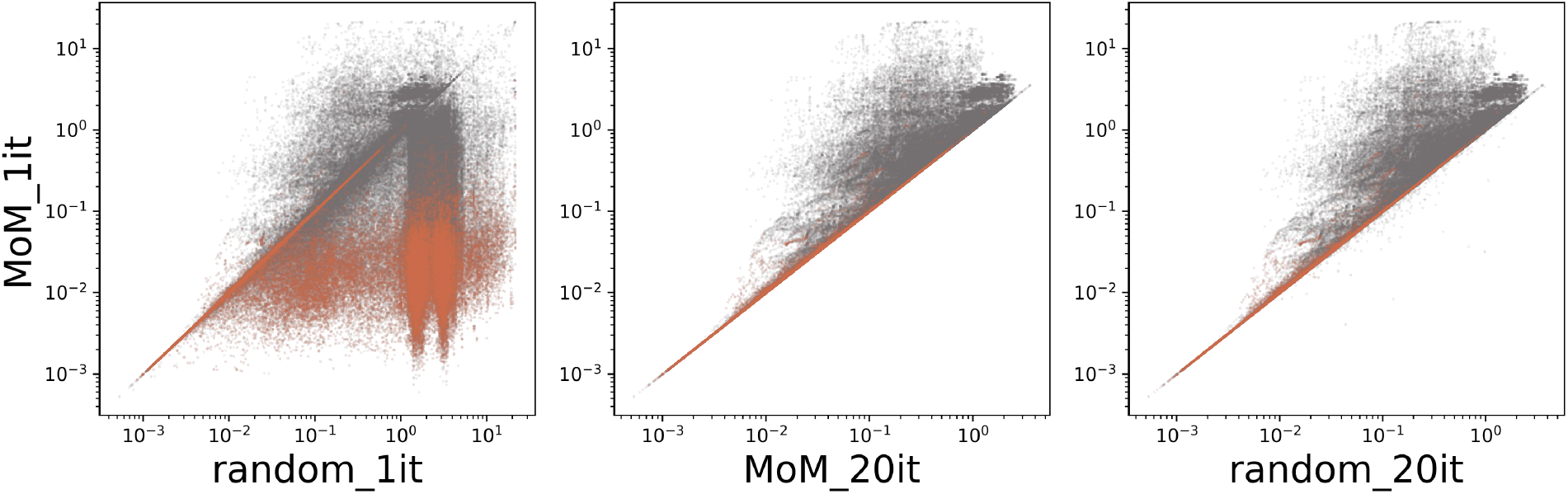
Performance of the standard search procedure (method of moments initialization, 1 iteration of gradient descent) benchmarked against three alternatives with random initialization and 20-iteration searches. Value indicates the magnitude of the KL divergence at search termination (orange: genes at grid points in lowest quartile of total divergence, computed from `MoM_20it`; gray: genes at grid points outside).

#### S6.2 Results

**Figure S4:**
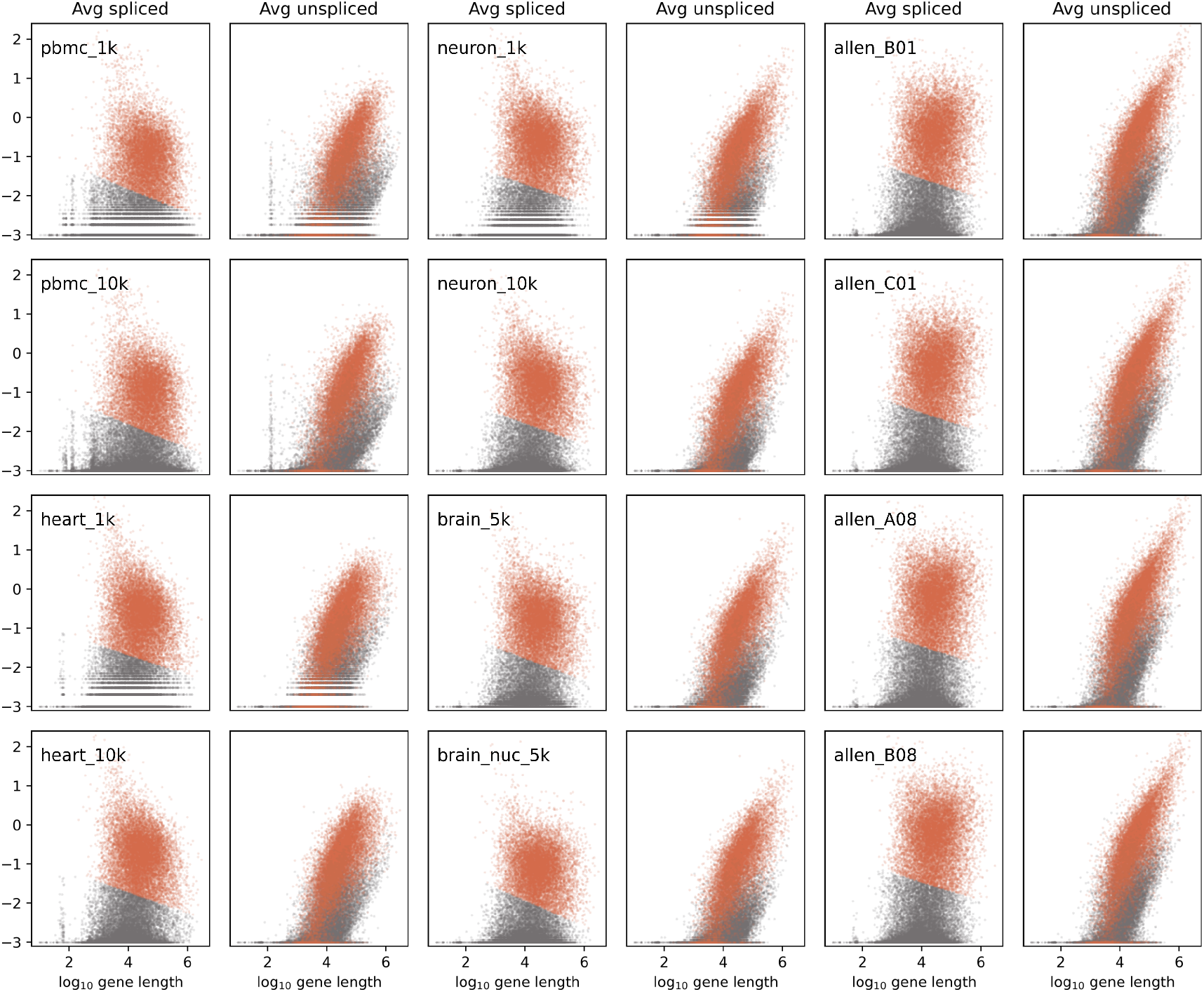
Length dependence of average spliced and unspliced mRNA observations in twelve datasets (orange: high-expression gene cluster; gray: discarded low-expression cluster). All datasets show overrepresentation of long unspliced mRNA, as well as separation into distinct high- and low-expression clusters.

**Figure S5:**
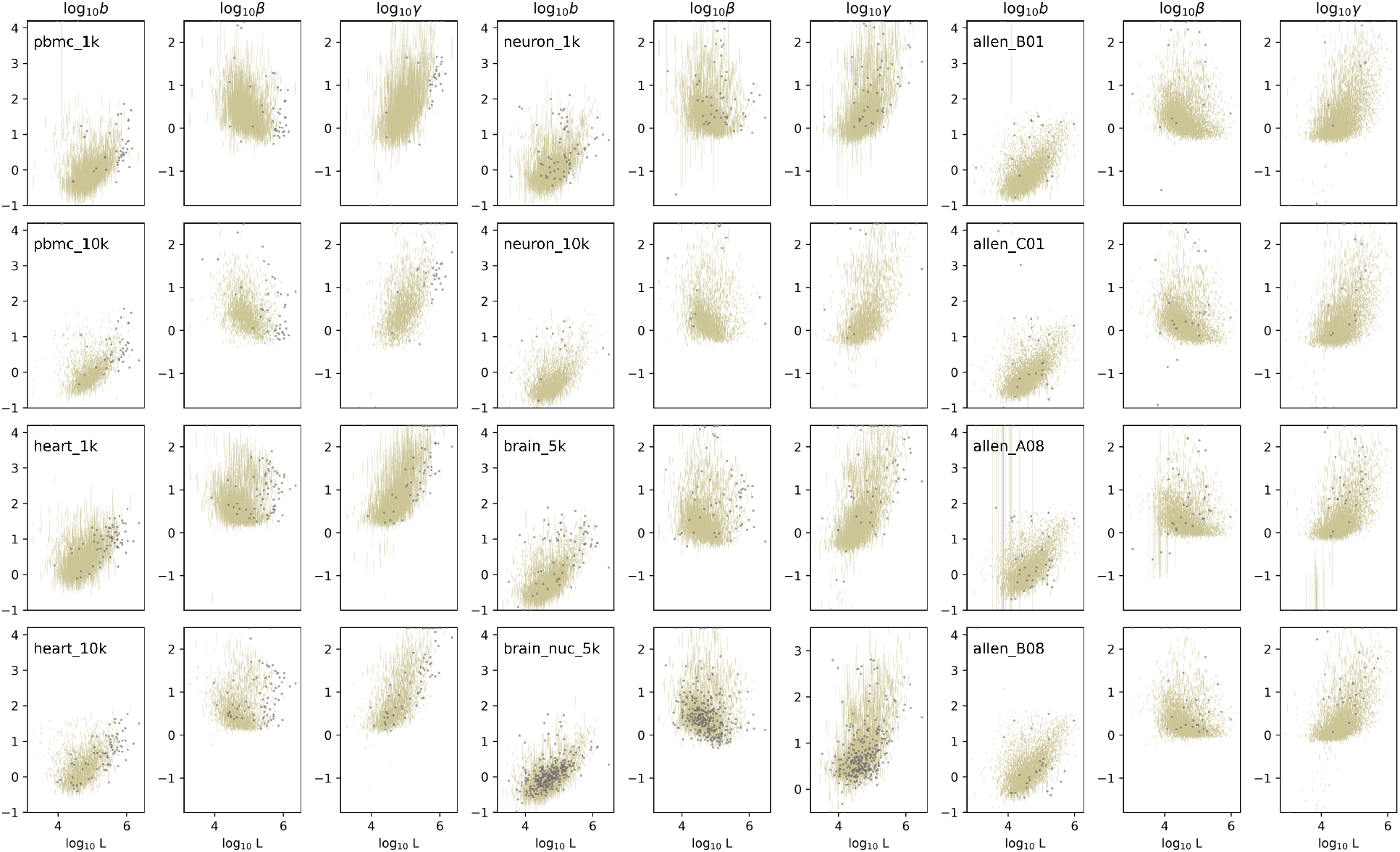
Transcriptional parameter estimates without a stochastic model of sequencing demonstrate pervasive length-dependent trends throughout all datasets (gold: lower bounds on 99% confidence intervals; gray: fits rejected by statistical testing).

**Figure S6:**
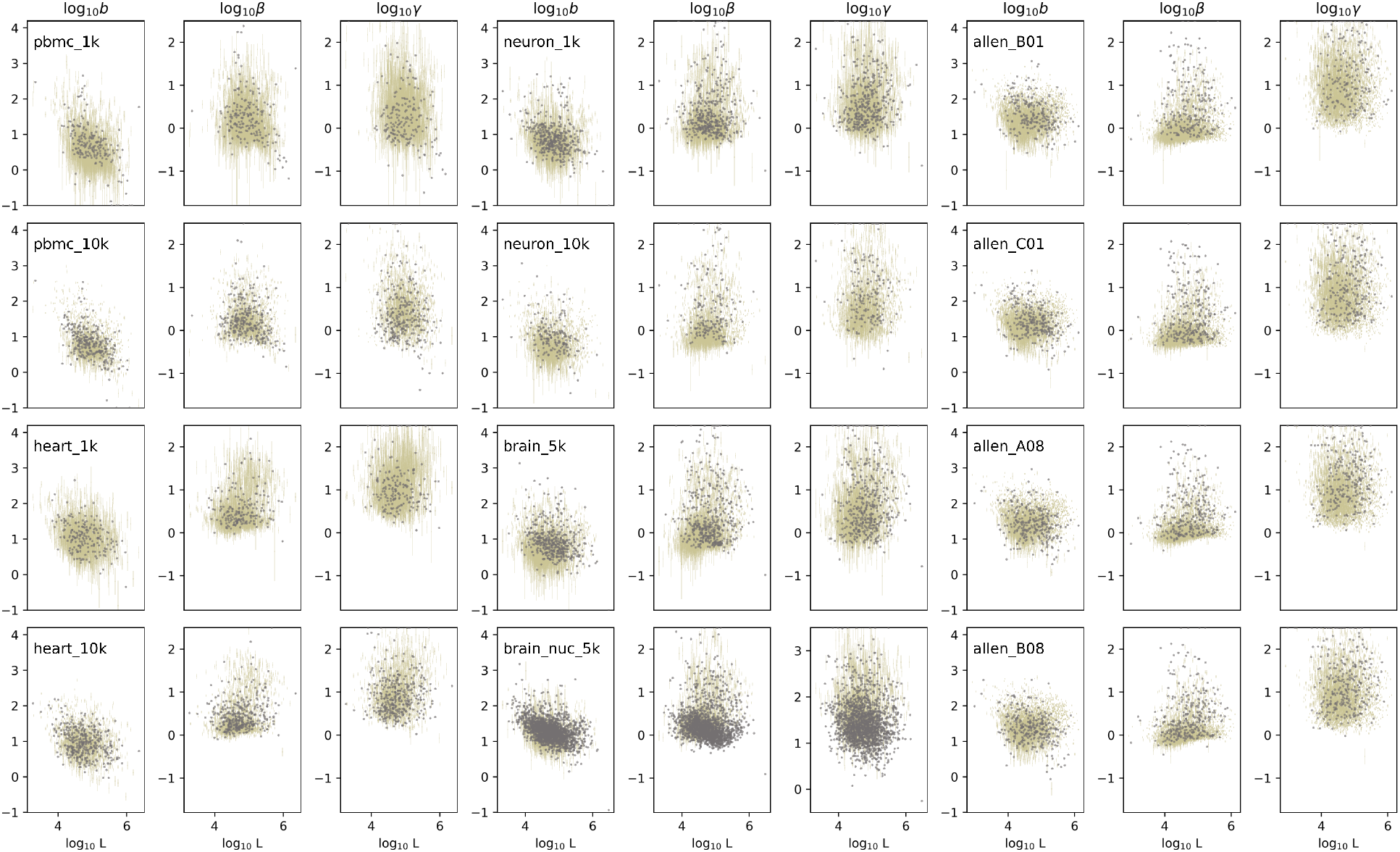
Inferred transcriptional parameters for the stochastic model of sequencing do not appear to have strong length dependence in any dataset (gold: lower bounds on 99% confidence intervals; gray: fits rejected by statistical testing).

**Figure S7:**
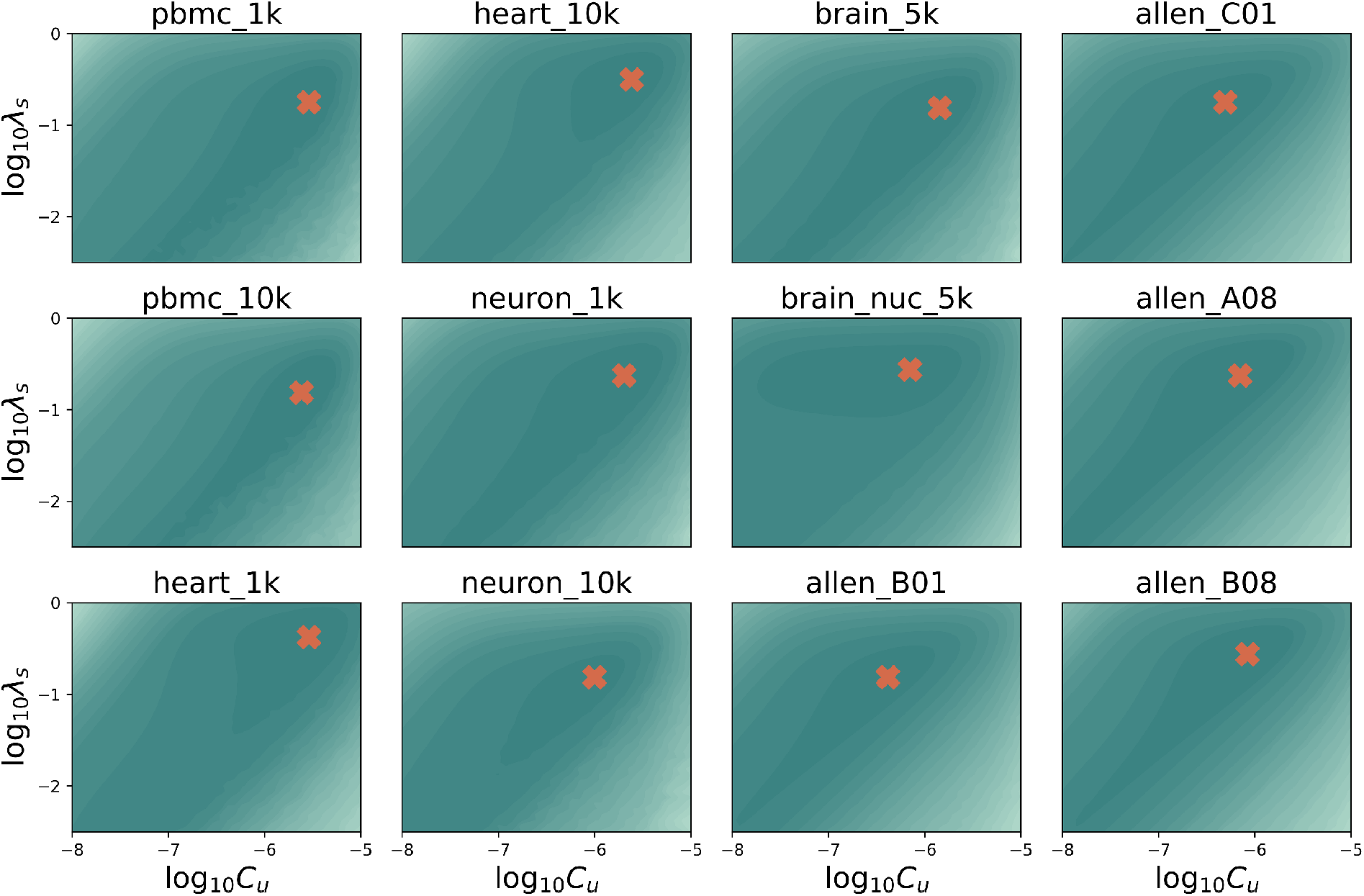
The sampling parameter likelihood landscapes show a single optimum in each dataset (dark: lower, light: higher total Kullback-Leibler divergence between fit and data; orange cross: optimal sampling parameter fit).

**Figure S8:**
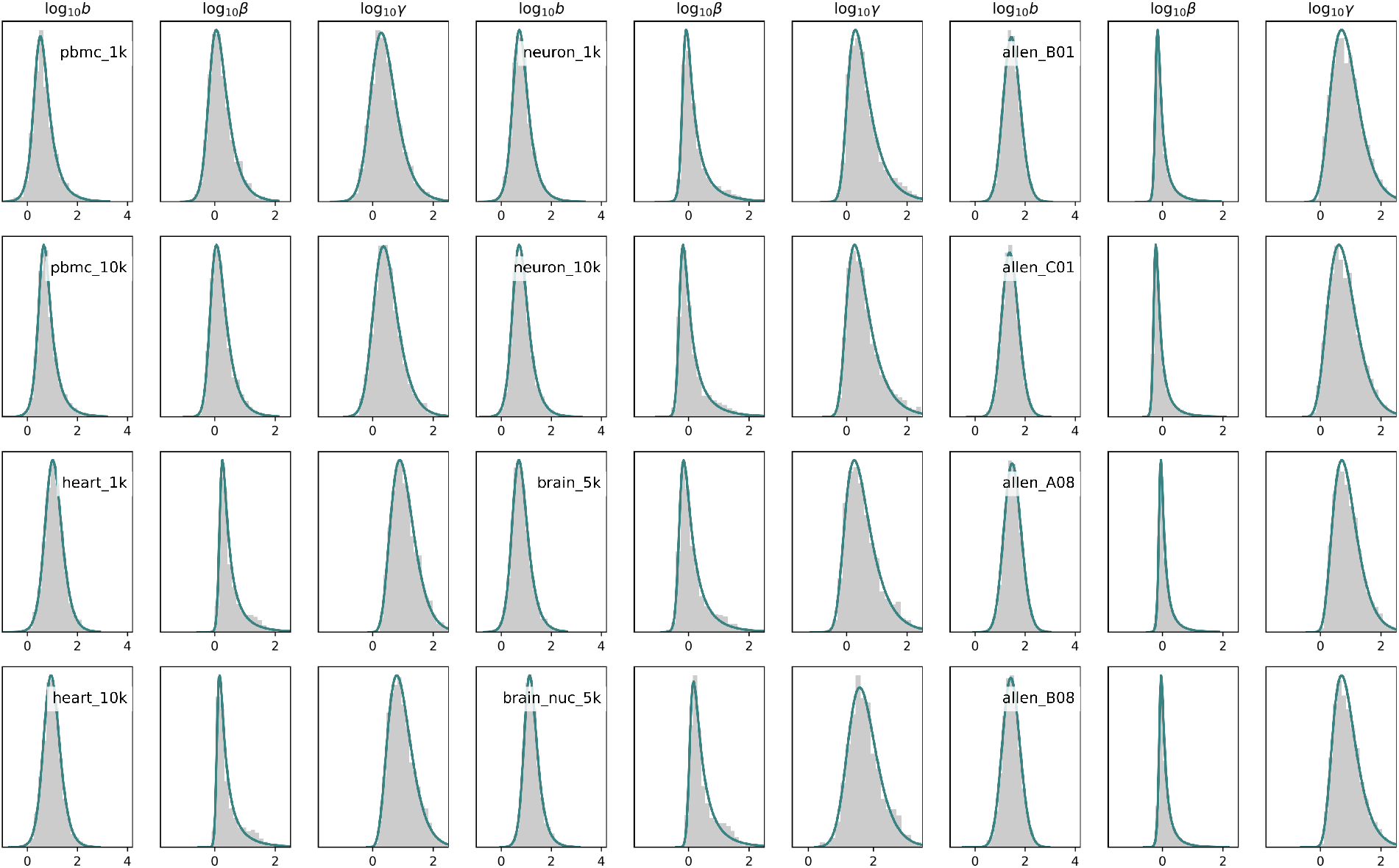
Inferred transcriptional parameter distributions are well-described by an normal-inverse Gaussian fit (gray: histogram of genes retained after statistical testing; teal line: best fit to normal-inverse Gaussian distribution).

**Figure S9:**
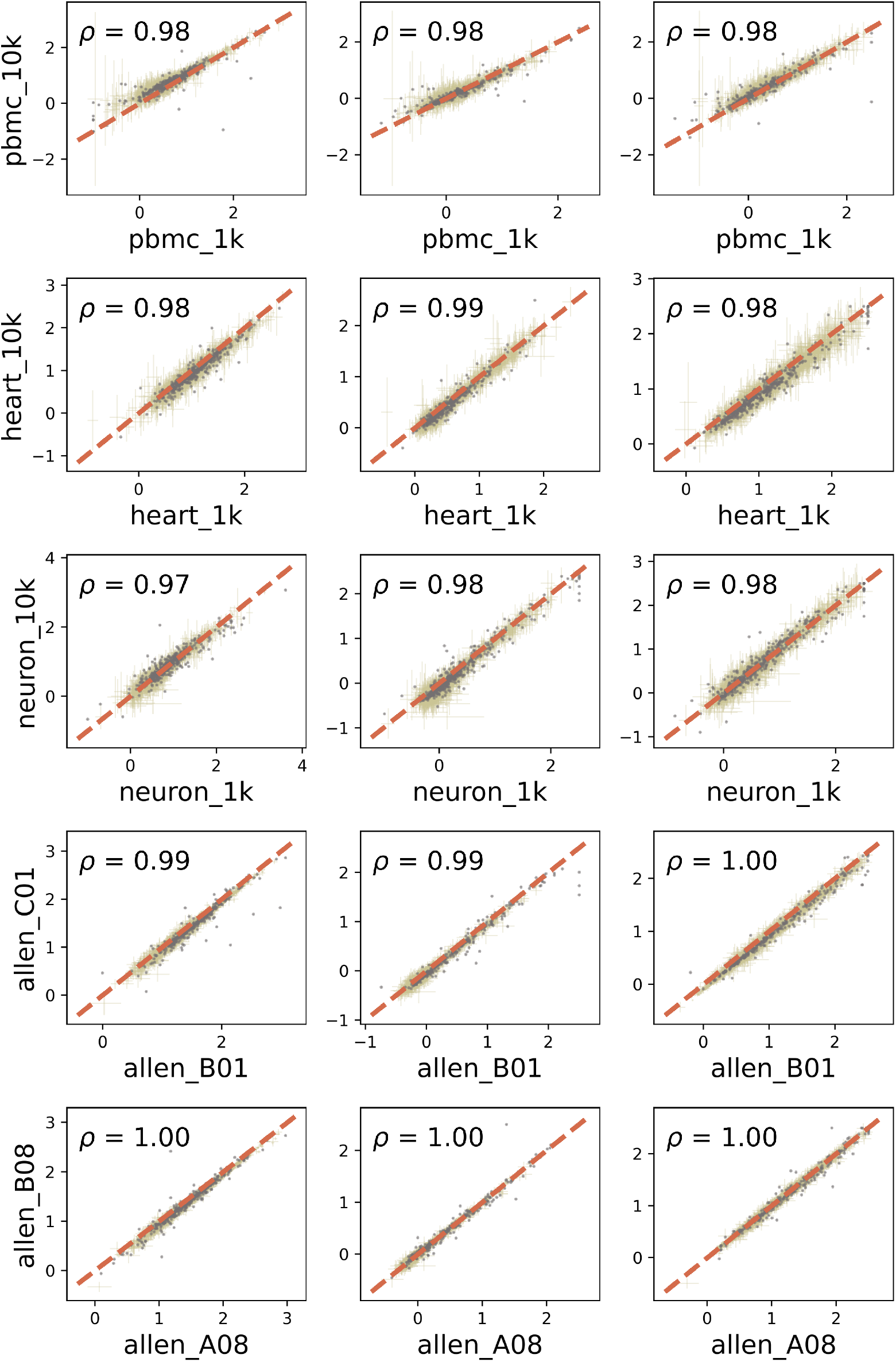
All five technical replicates show largely concordant inferred parameter values (orange dashed line: identity; gold: lower bounds on 99% confidence intervals; gray: fits rejected by statistical testing).

**Figure S10:**
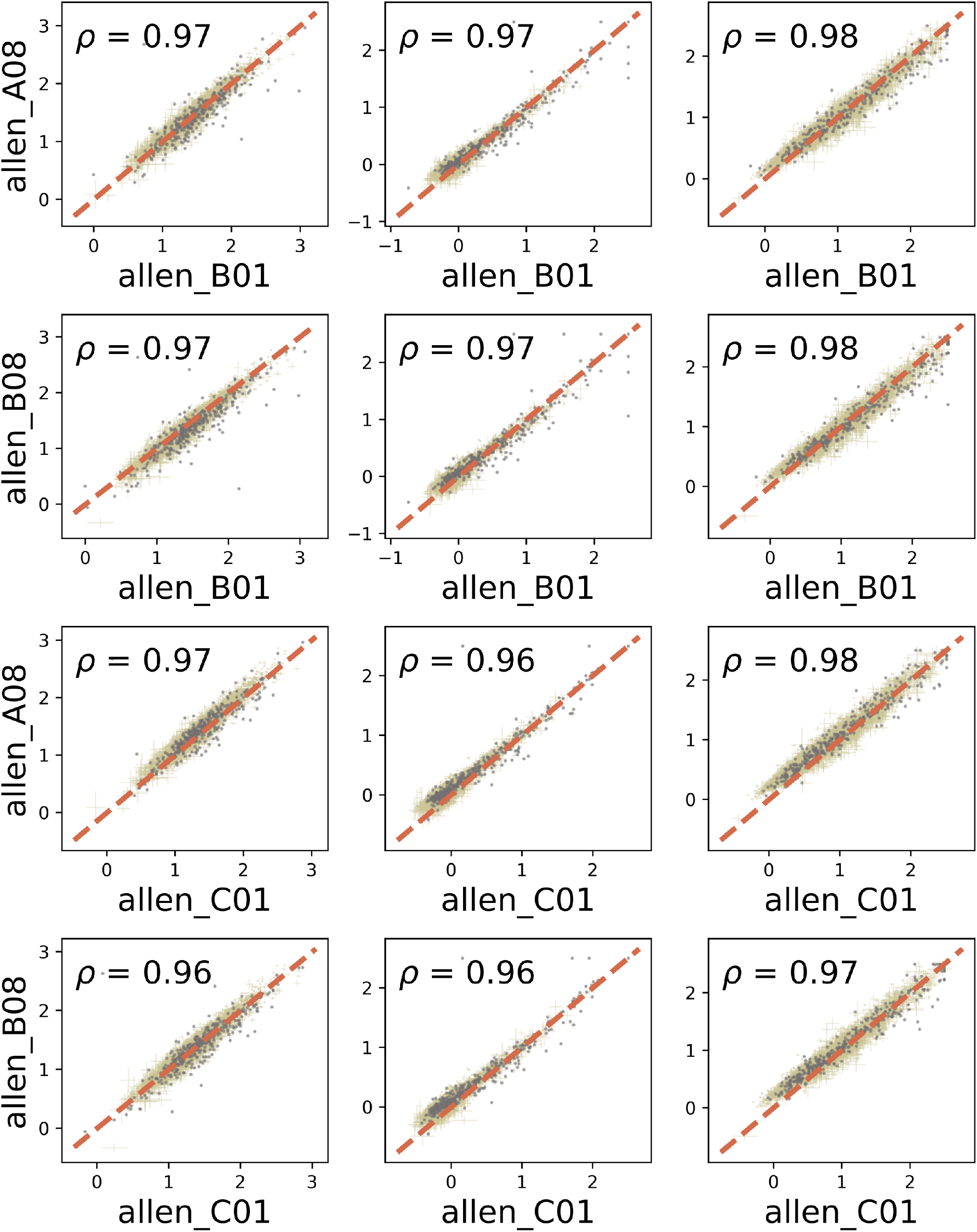
All four biological replicates show largely concordant inferred parameter values, albeit with lower correlations than technical replicates (orange dashed line: identity; gold: lower bounds on 99% confidence intervals; gray: fits rejected by statistical testing).

**Figure S11:**
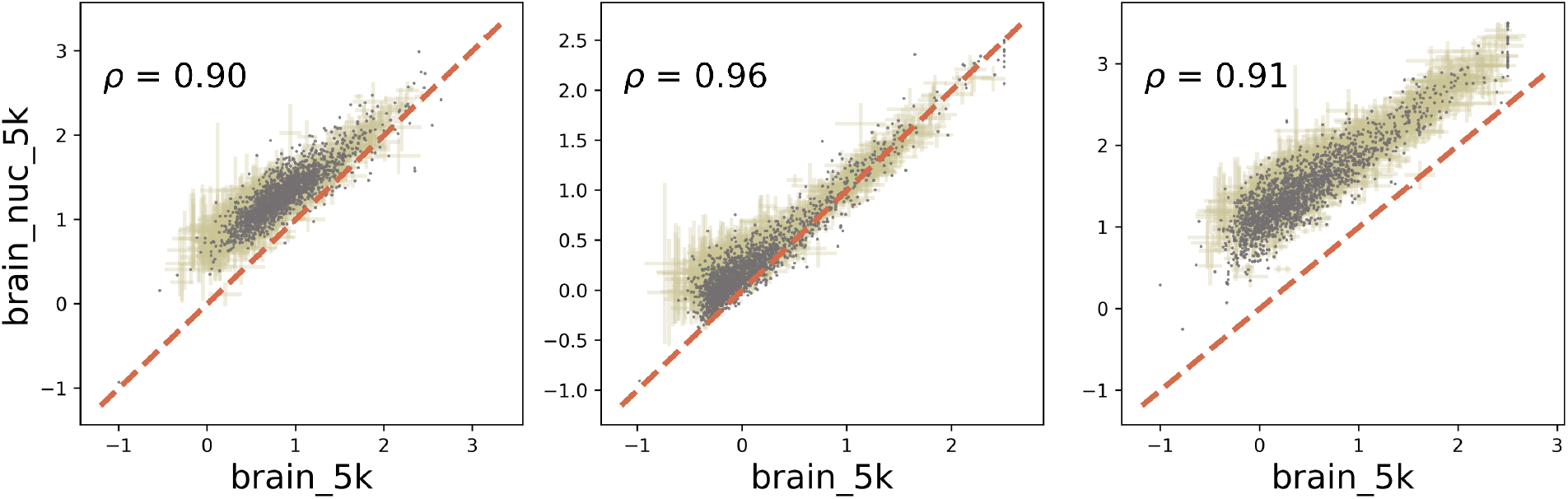
Nuclear/whole-cell scRNA-seq replicates appear to be qualitatively concordant. However, the nuclear RNA dataset shows a high rate of rejection and “degradation” parameters approximately one order of magnitude higher than those in the matched whole-cell dataset (orange dashed line: identity; gold: lower bounds on 99% confidence intervals; gray: fits rejected by statistical testing).

**Figure S12:**
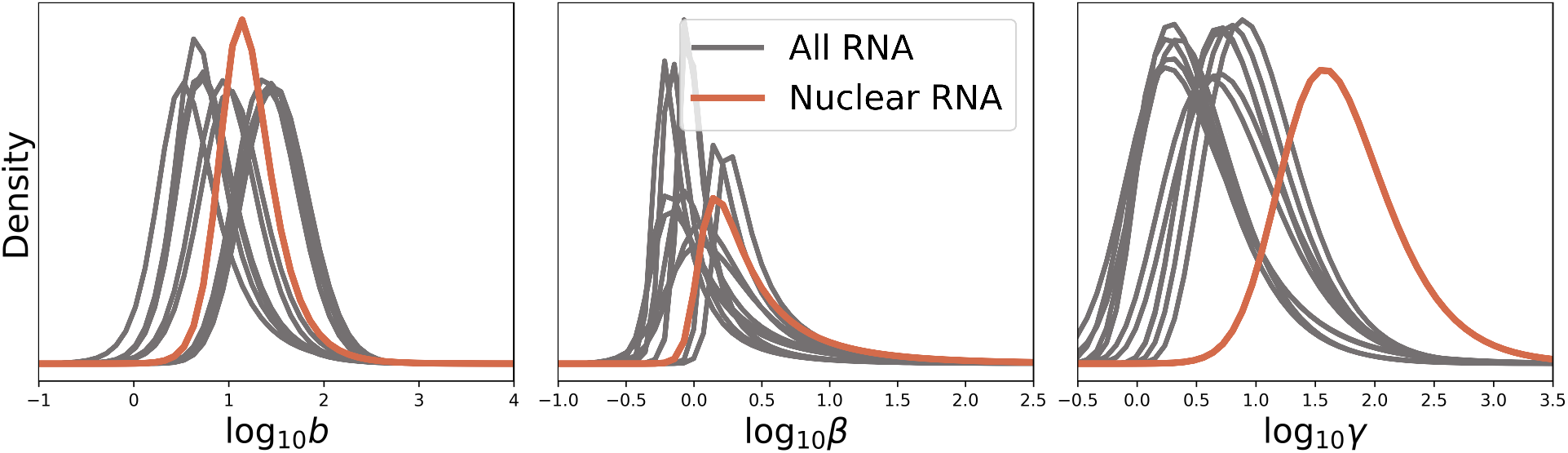
A comparison of all normal-inverse Gaussian fits to inferred parameter distributions, as reported in Fig. S8. As suggested by the offset in the matched dataset comparison (Fig. S11), the nuclear RNA have an average “degradation” or efflux rate 1-2 orders of magnitude higher than the other, whole-cell datasets.

**Figure S13:**
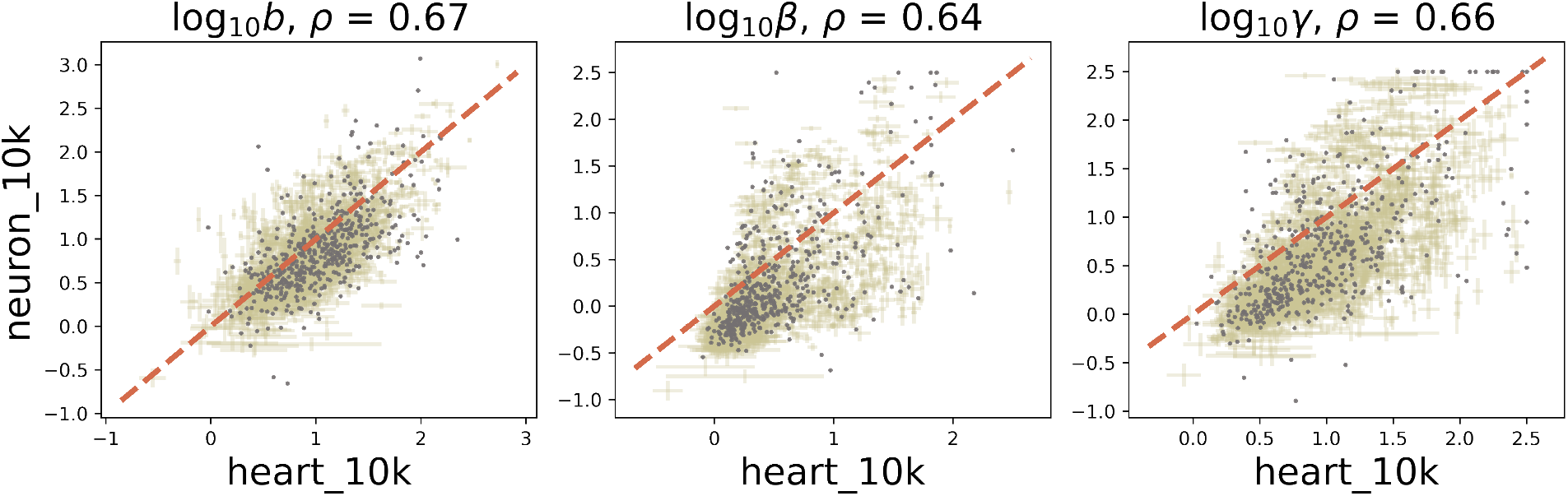
A comparison of parameter fits to 10X mouse heart and brain datasets. The parameters are correlated, but show significant tissue-specific deviations unobserved in biological and technical replicates. An offset from the identity line is evident, possibly due to suboptimal sampling parameter fits (orange dashed line: identity; gold: lower bounds on 99% confidence intervals; gray: fits rejected by statistical testing).

**Figure S14.**
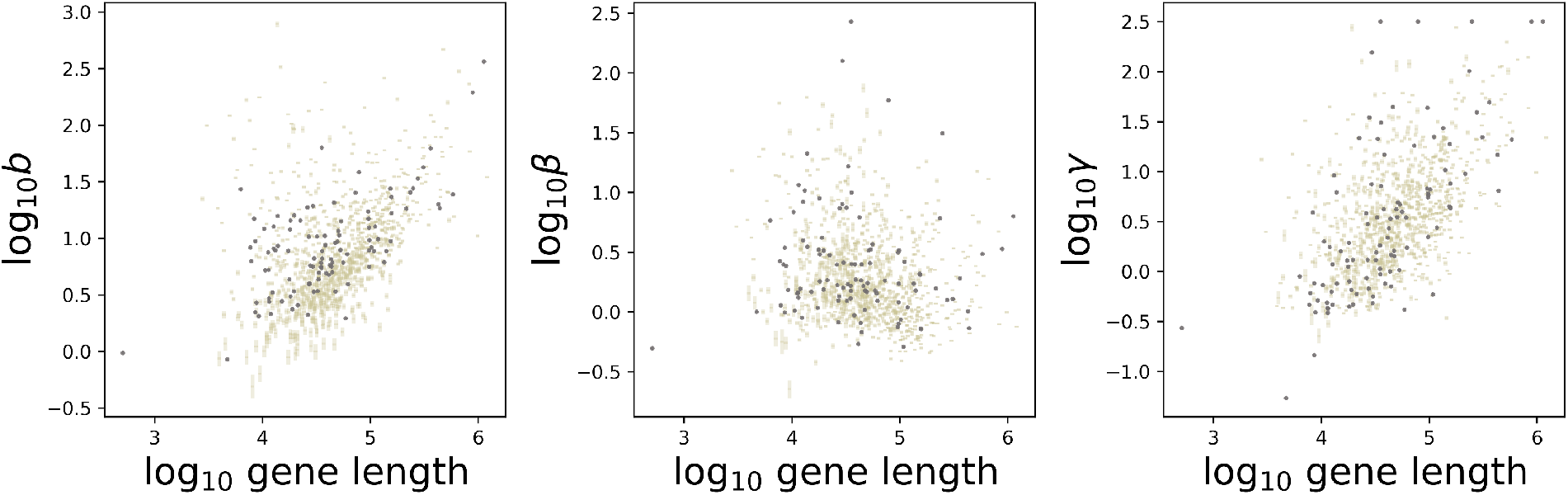
Parameter fits resulting from the Poisson, length-independent model of sequencing. The trends are qualitatively identical to those in Fig. S5, up to translation (gold: lower bounds on 99% confidence intervals; gray: fits rejected by statistical testing).

